# The coiled coil domain of DEF6 facilitates formation of large vesicle-like, cytoplasmic aggregates that trap the P-body marker DCP1 and exhibit prion-like features

**DOI:** 10.1101/538082

**Authors:** Huaitao Cheng, Fred Sablitzky

**Affiliations:** From the Department of Molecular Cell and Developmental Biology, School of Life Sciences, Queen’s Medical Centre, University of Nottingham, Nottingham NG7 2UH, UK

**Keywords:** DEF6, P-body, Coiled coil, ITAM motif, Cell stress, Prion-like

## Abstract

DEF6, also known as SLAT and IBP, is critical for the development of autoimmune disease and cancer. In T cells, DEF6 participates in TCR-mediated signalling determining T helper cell-mediated immune responses. In addition, DEF6 acts as a guanine nucleotide exchange factor for Rho GTPases facilitating F-actin assembly and stabilisation of the immunological synapse. However, DEF6 is also a component of mRNA processing bodies (P-bodies) linking it to mRNA metabolism. DEF6 can adopt multiple conformations that result in different cellular localisations and functions. Post translational modifications such as phosphorylation result in conformational change liberating functional domains that are masked in the native stage of DEF6. ITK phosphorylation of Try210/222 liberates the N-terminal end and to a certain extend also the C-terminal coiled coil domain of DEF6 resulting in P-body colocalisation. In fact, the N-terminal 45 amino acids of DEF6 that encode a Ca2+-binding EF hand are sufficient to target P-bodies. Mutant proteins that unleashed the C-terminal coiled coil domain of DEF6 spontaneously aggregated forming large vesicle-like, cytoplasmic structures. These aggregates trapped proteins such as the P-body component DCP1 altering its cytoplasmic localisation. However, cellular stress reversed aggregate formation in mutant DEF6 proteins that contained ITAM and PH domain in conjunction with the coiled coil domain resulting in colocalisation with DCP1. Furthermore, coiled coil-mediated aggregates appeared to function like prions enforcing conformational change onto wild type DEF6 protein.

---

DEF6 exhibits strong homology to only one other protein namely SWAP70 (1 to 5). Both proteins are highly conserved exhibiting the same domain structure and originate from a common ancestor gene present in *tricoplax adhaerens* (6). Two Ca 2+-binding EF hands at the N-terminus are followed by an immunoreceptor tyrosine-based activation motif like (ITAM) sequence. A central pleckstrin-homology (PH) domain, which acts as a PI (3,4,5) P3 anchor, is followed by the C-terminal Dbl homology-like (DHL) domain that contains a coiled coil domain and exhibits GEF activity (Fig. 1 A) (7, 8, 9).

**Figure 1.**
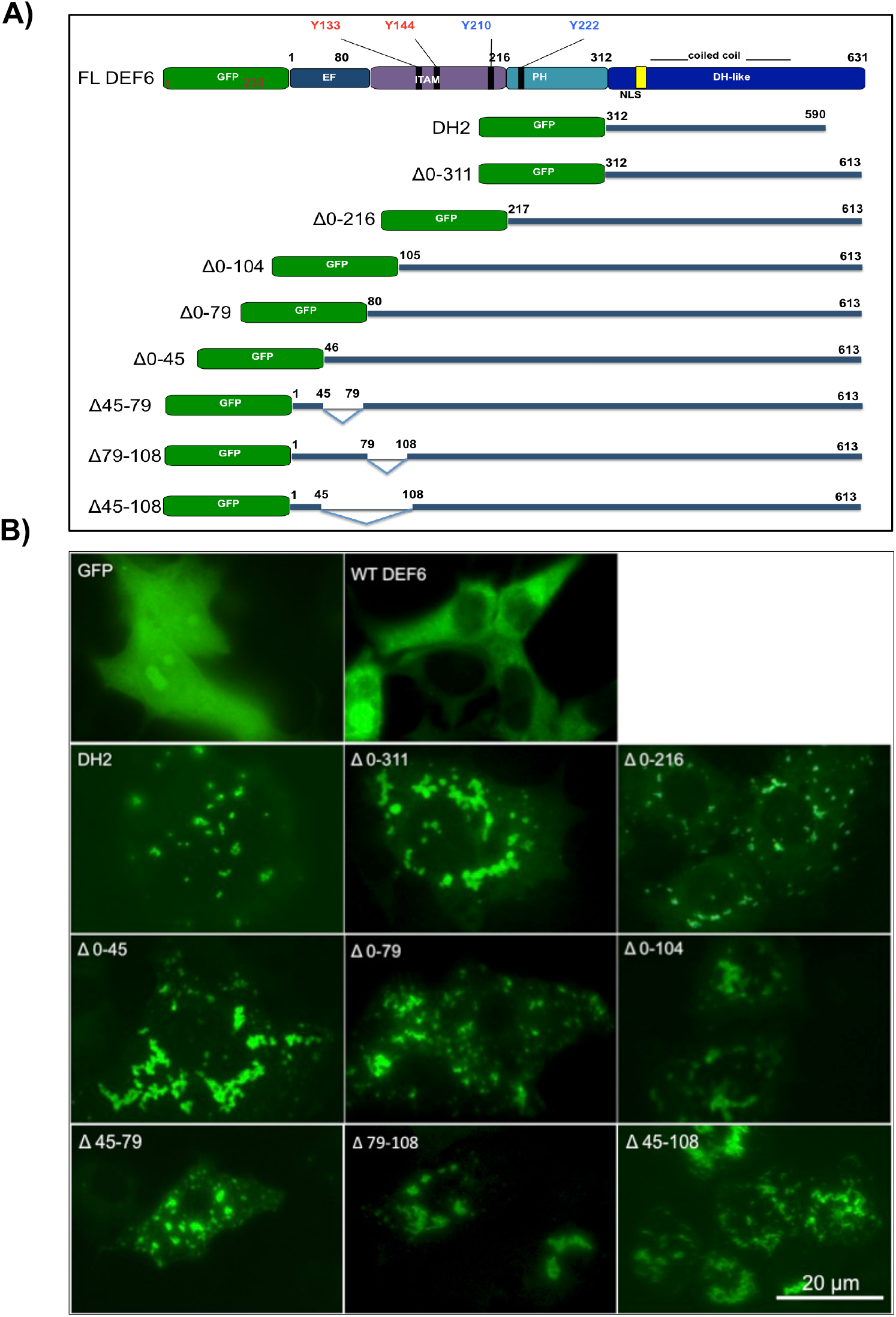
DEF6 N-terminal truncation and deletion mutants form large cytoplasmic aggregates. A) Schematic representation of DEF6 N-terminal truncation and deletion mutants. GFP-tagged N-terminal truncation and deletion mutants were established as indicated. B) 24h after transfection of COS7 cells, GFP-tagged N-terminal truncation and deletion mutants as indicated exhibited large cytoplasmic aggregations. GFP and wild type DEF6 are shown for comparison.

DEF6 structure and function is regulated through post-translational modifications. In T cells, two tyrosine kinases, LCK and ITK, phosphorylate DEF6 resulting in conformational change, cellular translocation and differential activity. Phosphorylation of tyrosine residues 133, 144 (8) and possibly 210 (10) by LCK resulted in recruitment of DEF6 to the immunological synapse (IS) and guanine nucleotide exchange (GEF) activity (8, 10). Phosphorylation of tyrosine residues 210 and 222 by ITK resulted in cytoplasmic granule formation that colocalise with DCP1, a component of P-bodies (9).

P-bodies are messenger ribonucleoprotein (mRNP) granules that facilitate mRNA 5’ to 3’ decay and RNA interference (RNAi) (11 to 14 and reviewed in 15). P-bodies are non-membranous dynamic supermolecular complexes, containing more than 34 types of proteins including DEF6 (9, 10, 13, 14, 16). Many P-body components contain coiled coil domains, such as Edc4, Pdc1 and mLin41 (17, 18). Decker and Parker (2012) summarised a possible model of P-body assembly in yeast, which depends on EDC3 self-interaction domain (Yjef-N) and Lsm4 Q/N rich coiled coil domain (19). Although in Drosophila S2 cells, P-body assembly is not blocked by EDC3 depletion, to deplete some P-bodies Q/N rich proteins, such as GW182, would decrease P-body formation in Drosophila and human cells. In fission yeast *Schizosaccharomyces pombe*, Pdc1 interact with Dcp1 in P-bodies. A truncated mutant Pdc1 protein lacking its coiled coil region, failed to localise into P-bodies and P-body assembly was impaired (17). Similarly, deletion of the coiled coil domain of mLin41 resulted in reduced interaction between mLin4 and Ago2 in Hela cells (18). It is likely therefore that assembly through interaction of coiled coil domains is an important step for P-body formation and function (17, 18, 20).

The coiled coil domain is a super-secondary protein structure in which two to seven alpha helices interact with each other forming a super-coiled structures (21). Coiled coil domains are found in 3% to 5% of all proteins and have been frequently been shown to mediate protein-protein interactions (22, 23, 24).

Ectopic expression of wild type DEF6 protein tagged with GFP resulted in diffuse cytoplasmic localisation but DEF6 ITK phosphomimic mutant, DEF6-Y210E/222E, overexpressed in COS7 cells exhibited a different cellular localisation forming DEF6 foci that overlapped with DCP1 (9). Cellular stress induced through arsenite treatment resulted in granule formation of wild type DEF6 that colocalised with DCP1 (9). Similarly, endogenous DEF6 in Jurkat cells has also been observed to colocalise with P-bodies (9, 25).

To further elucidate the function of the coiled coil domain of DEF6, a comprehensive analysis of mutant DEF6 proteins overexpressed as GFP-fusion proteins in COS7 cells was conducted. Unleashing the coiled coil domain triggered formation of large, vesicle-like aggregates of the mutant DEF6 proteins. These aggregates did not overlap with mCherry-tagged DCP1 but instead trapped DCP1 altering its normal cellular distribution. Cellular stress through treatment with either arsenate or nocodazole reversed aggregation of mutant DEF6 proteins that contained the ITAM and PH domain in addition to the coiled coil domain, resulting on colocalisation with DCP1. Furthermore, coiled coil-mediated aggregates induced aggregation of wild type DEF6 protein suggesting that the coiled coil domain of DEF6 can enforce conformational change in a prionlike fashion.

## RESULTS

### The N-terminal end of DEF6 restrains the C-terminal coiled coil domain from formation of large cytoplasmic aggregations

We have shown that the N-terminal end of DEF6 is sufficient to spontaneously colocalise with the P-body marker DCP1 whereas the C-terminal coiled coil domain was dispensable (26). To establish the functional role of the coiled coil domain, N-terminal truncation and deletion mutants were established (Fig. 1A) and expressed in COS7 cells (Fig. 1 B). Remarkably, all 9 mutant DEF6 proteins formed cytoplasmic aggregates regardless of which of the N-terminal amino acids were missing (Fig. 1B). In particular, DH2 and Δ0-311 that contained just the C-terminal end including the coiled coil domain formed large aggregates suggesting that the N-terminal domains of DEF6 in its wild type conformation restrain the coiled coil domain from aggregation.

To further test the role of the coiled coil domain in aggregation formation, a series of DEF6 truncation mutants were established, which either contained the coiled coil domain entirely or partially or lacked it completely as indicated in Fig. 2A. As shown in Fig. 2B, DH2 that just contained the coiled coil domain formed aggregates. In contrast, DH1 and DHL-C (partially containing the coiled coil domain) did not form aggregates. Instead, these mutant proteins labelled lamellipodia, filopodia and stress fibres (Fig. 2B). Mutant DEF6 proteins not containing the coiled coil domain (ITAM, PH1 and DHL-N) localised diffuse in cytoplasm and nucleus without any signs of aggregations. Nuclear localisation of ITAM and PH1 mutant are likely due to their small size. In contrast, PH2 which contains a predicted NLS region (amino acid positions 330 to 341) localised exclusively to the nucleus as previously shown (27). DHL-N also partially contains the coiled coil region. As shown in Figure 2B, this mutant protein was mainly distributed diffuse in the cytoplasm and was absent from the nucleus. However, DHL-N did occasionally exhibit distinct cytoplasmic granules. These granules have three features: 1) only one or two are seen per cells; 2) they always localise next to the cell nucleus; 3) if present as a pair, they usually localised on the opposite sides of the cell nucleus and it remains to be seen whether this localisation of DHL-N coincides with centrosomes.

**Figure 2.**
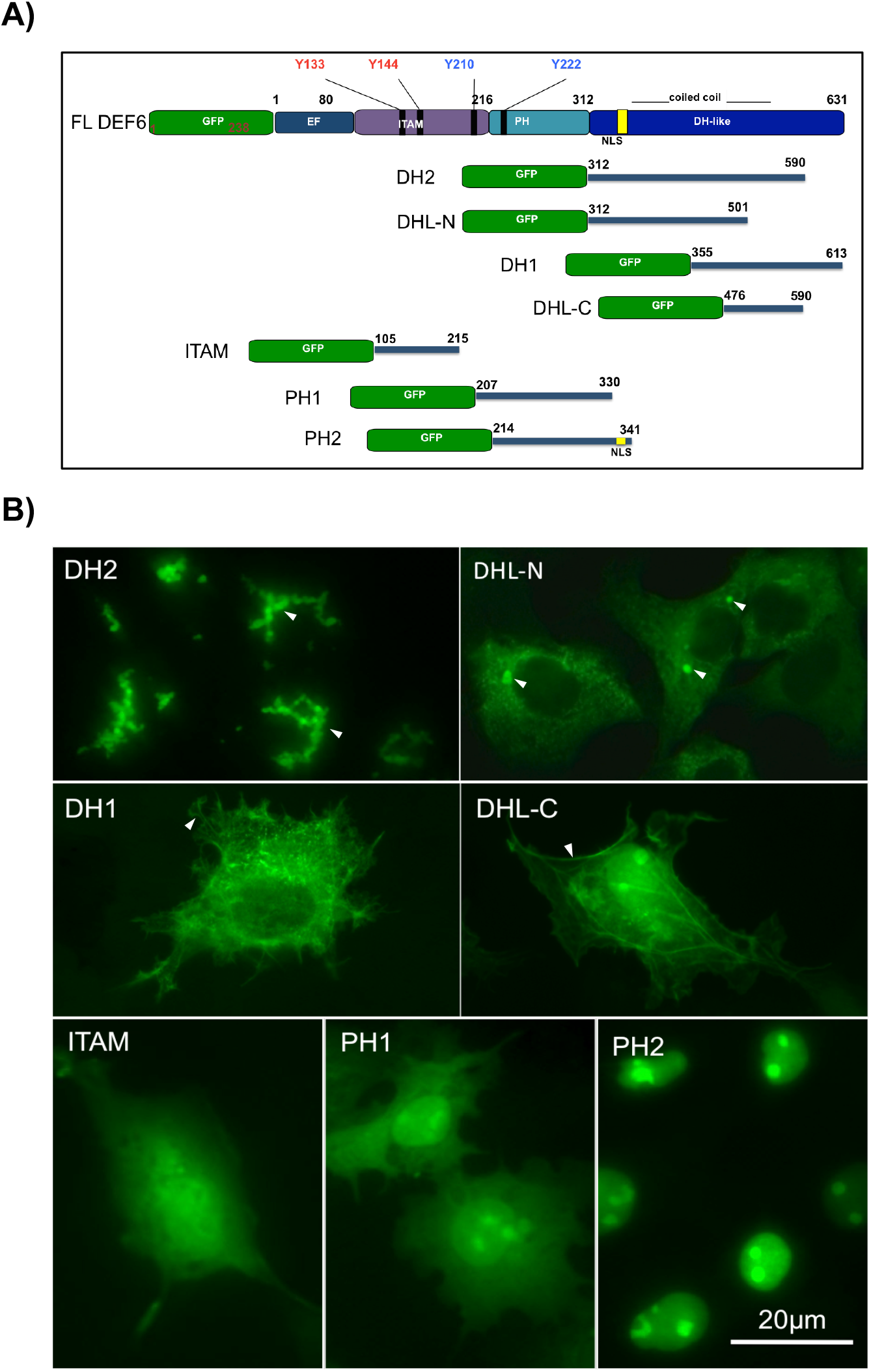
The coiled coil domain of DEF6 facilitates aggregation. A) Schematic representation of DEF6 N-terminal truncation and deletion mutants. B) GFP-tagged mutants schematically depicted in Fig. 3.7.2a were transfected into COS7 cells and images taken 24h after transfection. DH2 that contains the entire coiled coil region formed large aggregates in the cytoplasm. DHL-N that partially contains the coiled coil region was mainly diffuse in cytoplasm. However, some large round granules were occasionally observed with the following features: only one or two granules were present in the cytoplasm that always localised next to and when present in pairs at opposite sides of the cell nucleus. DH1 and DHL-C did not form large aggregates despite containing the C-terminal part of the coiled coil domain. Instead, they localised to lamellipodia, fílopodia and stress fibres. The ITAM and PH1 domain tagged to GFP were diffuse in cytoplasm and nucleus similar to GFP alone. The PH2 mutant that contained the predicted NLS sequence of DEF6 was exclusively localised in the nucleus as previously shown (27).

To firmly establish that the coiled coil domain is indeed facilitating DEF6 aggregation, mutants Δ0-104-All-10 and Δ0-216-All-10 were established and tested (Fig. 3). In these mutants the coiled coil domain was disrupted by the introduction of proline residues in 10 highly conserved a/d positions as described in Cheng et al. (26). The N-terminal truncation mutants Δ0-104 and Δ0-216 formed consistently large aggregations, whereas Δ0-104-All10 and Δ0-216-All-10 did not. Instead, these two coiled coil disrupted mutants exhibited similar phenotype to the All-10 mutant (Fig. 3) (26) localising to lamellipodia and filopodia. Together the data established that the coiled coil domain facilitates DEF6 aggregation and this function is masked/inhibited in the wild type protein.

**Figure 3.**
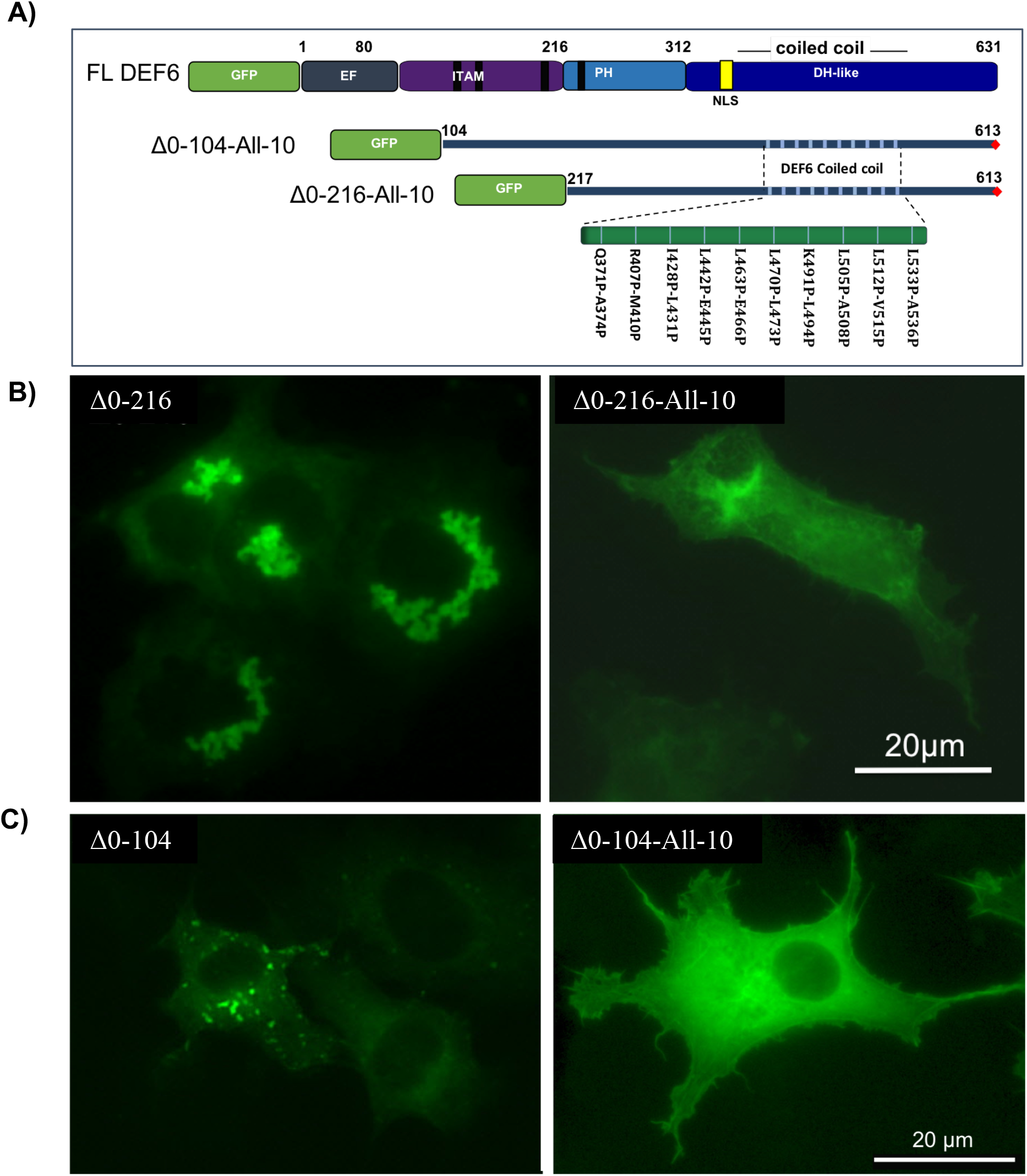
Coiled coil-mediated formation of large cytoplasmic aggregates is abolished through proline mutations that disrupt the coiled coil domain of DEF6. A) Schematic representation of GFP-tagged Δ0-104-A11-10 and Δ0-216-All-10 DEF6 mutants. 24 h after transfection of COS7 cells, GFP-tagged Δ0-216 and Δ0-104 mutant formed aggregates that seemed to interact with each other forming large structures (B and C left panel). Aggregation and large structure formation was abolished when the coiled coil domain was disrupted through prolines at 10 a/d positions (B and C right panel).

### Coiled coil-mediated aggregates of DEF6 form large structures that ‘trap’ the P-body marker DCP1

To further test the functional properties of DEF6 aggregates, cotransfection with DCP1 were performed. As shown in Figures 4, DEF6 aggregates differed in their shape and structure but were always associated with DCP1 albeit in various ways clearly distinct from the colocalisation with DCP1 described above. (i) DH2 aggregates were unique in the sense that they formed vesicle-like tubular structures (Fig. 4 I). DCP1 was always associated with these aggregates but never colocalised. Furthermore, cellular localisation of DCP1 was clearly altered: the normal cytoplasmic distribution of DCP1/P-bodies was absent from the double transfected COS7 cells and all DCP1 was tightly ‘bound’ by the DEF6 aggregates. In addition, the normally round shape of DCP1/P-bodies was altered exhibiting shapes that appeared to be dependent upon the shape of DEF6 aggregates (Fig. 4). DH2 vesicle-like tubular aggregates somewhat resembled the structure of the Golgi apparatus but cotransfection of DH2 with several Golgi markers (Addgene: mCherry-GalT and mCherry-SiT-N-15) were inconclusive (data not shown) and it remains to be seen whether DH2 aggregates are associated with some cytoplasmic organelles or self-assembled. (ii) Δ0-216 and Δ0-311 did not form vesicle-like aggregates as seen with DH2, but were also always associated with DCP1 altering its cytoplasmic localisation (Fig. 4 II and Suppl. Fig. 3). (iii) All other mutant proteins tested fall into the 3rd group (e.g. Δ0-104): their aggregates were also not vesicle-like but did partially overlapped with DCP1 (Fig. 4 III and Suppl. Fig. 4 to 8).

**Figure 4.**
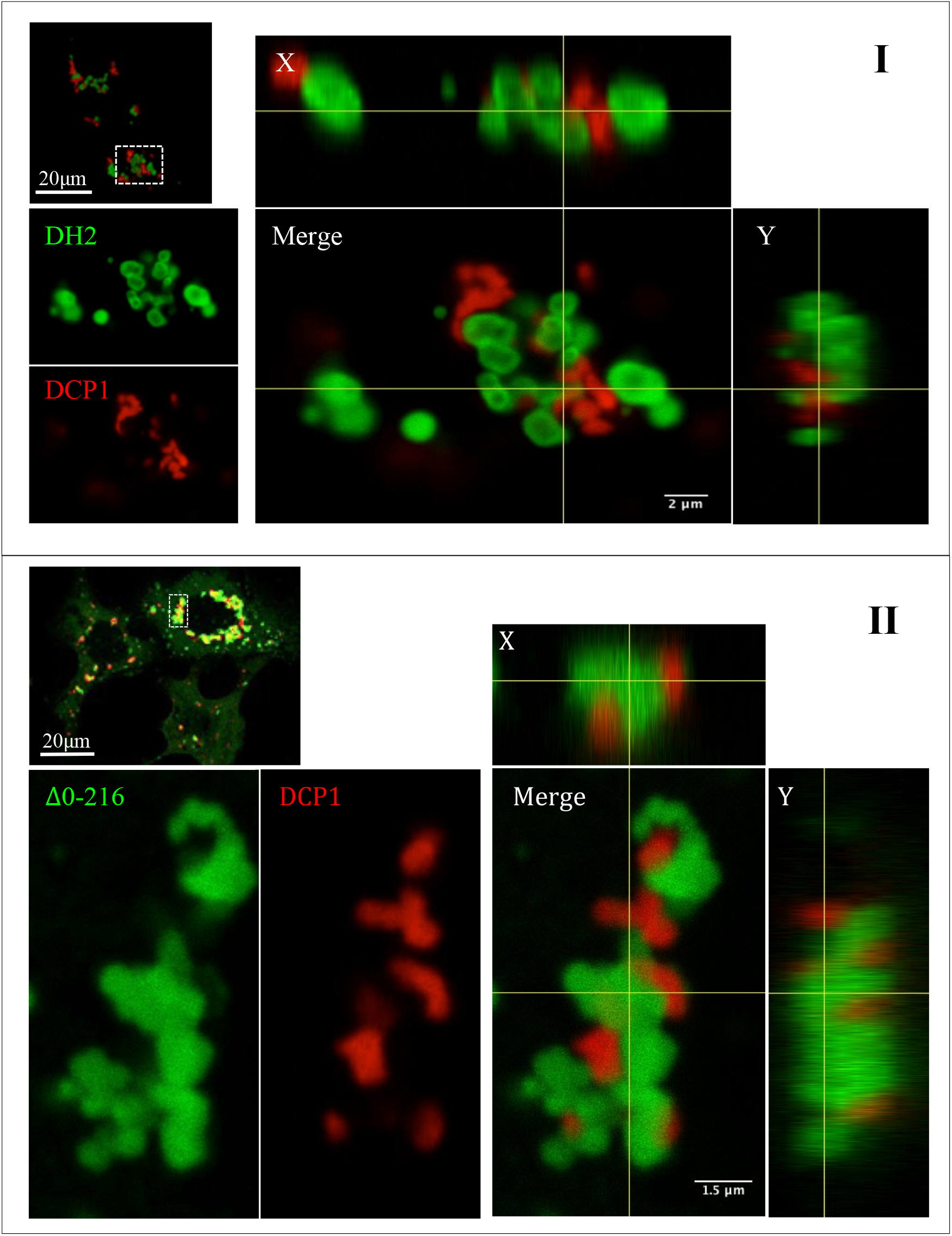
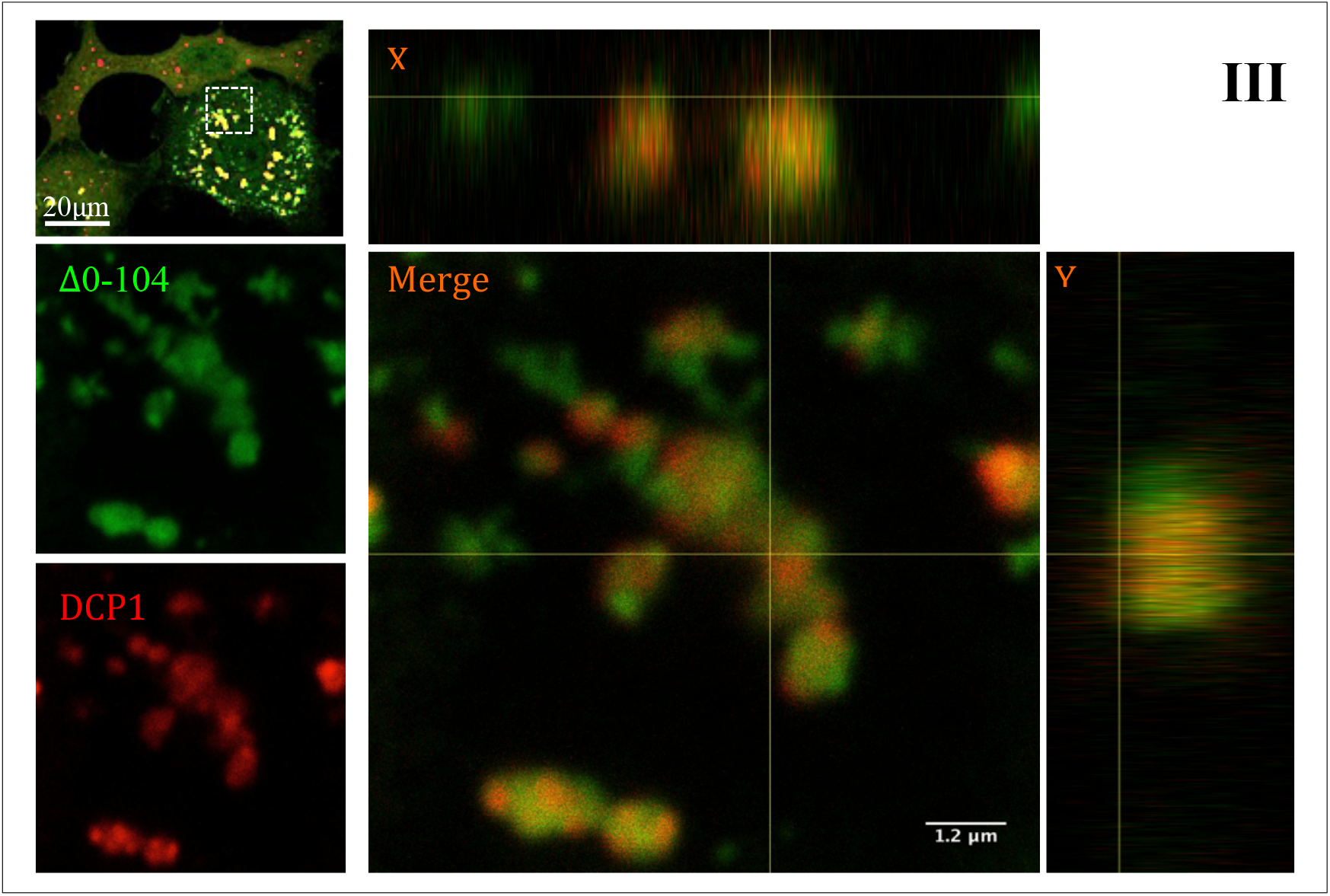
Coiled coil-mediated aggregates of DEF6 form large structure that ‘trap’ DCP1. Confocal analysis of transfected COS7 cells as described before. (I) GFP-tagged DH2 aggregates formed vesicle-like structures that did not colocalise with DCP1. Instead, localisation of DCP1 was altered and always adjacent to DEF6 structures (see merged images on the right). (II) GFP-tagged Δ0-216 and Δ0-311 (Suppl. Fig. 3) aggregates also formed large structures but these did not appear vesicle-like. However, these structures also altered the localisation of DCP1 that was always associated with DEF6 structures (see merged images on the right). (III) GFP-tagged Δ0-104 is representative for all other N-terminal truncation mutants tested. GFP-tagged Δ0-104 formed aggregates and larger structures but in this case these structures partially overlapped with DCP1; again altering the normal cellular localisation of DCP1 (see merged images on the right).

### Differential response of DEF6 aggregates to cellular stress

Arsenate treatment of COS7 cells caused wild type DEF6 to relocate colocalising with P-bodies (9). To test other stress conditions, cells were treated with nocodazole, a microtubule inhibitor. As shown in Figure 5, wild type DEF6 formed granules that colocalised with DCP1 in response to nocodazole treatment similar to arsenate treatment. This is consistent with the studies showing that inhibiting microtubules promote P-body formation (28, 29) and might suggest that cellular relocalisation of DEF6 into P-bodies is also independent of microtubules.

**Figure 5.**
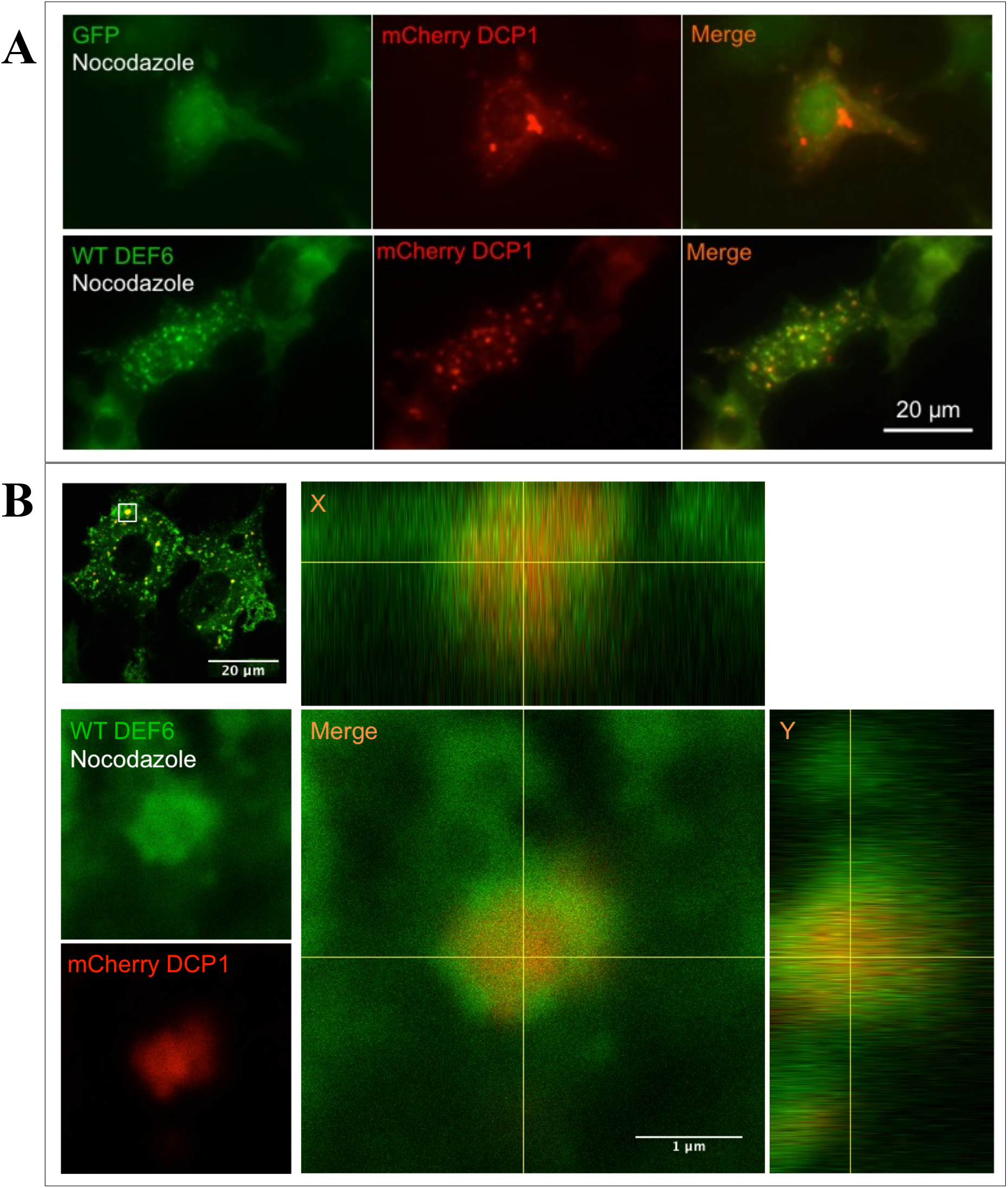
Cellular stress induced through nocodazole treatment results in WT DEF6 colocalising with DCP1. Fluorescent (A) and Confocal (B) analysis of COS7 cells expressing GFP-tagged wild type DEF6 and mCherry-tagged DCP1 after nocodazole treatment as described before. Merged images including vertical and side views (X and Y coordinates) shown on the right indicated colocalisation.

Coiled coil-mediated DEF6 aggregates responded to cellular stress in distinct fashion. Neither arsenate nor nocodazol treatment influenced aggregates formed by DH2(group 1) or mutant proteins of group 2 like Δ0-216 and Δ0-311. They still were associated with DCP1 as observed in untreated cells and never colocalised with DCP1 (Fig. 6 and 7 and Suppl. Fig. 9 and 12). In contrast, aggregates of mutant proteins of the 3rd group (e.g. Δ0-104) that partially overlapped with DCP1 in untreated cells mostly colocalised with DCP1 when treated with arsenate (Fig. 8 A and Suppl. Fig. 10 and 11) and completely colocalised with DCP1 after nocodazol treatment (Fig. 8 B Suppl. Fig. 13). It seems therefore that coiled coil-mediated aggregates that are formed by mutant DEF6 proteins either just containing the coiled coil domain (DH2) or in addition the PH domain (Δ0-216) are non-responsive to cellular stress. In contrast, in the presence of the ITAM domain (Δ0-104), aggregates partially overlap with DCP1 and can be induced to completely colocalise with DCP1 under stress conditions.

**Figure 6.**
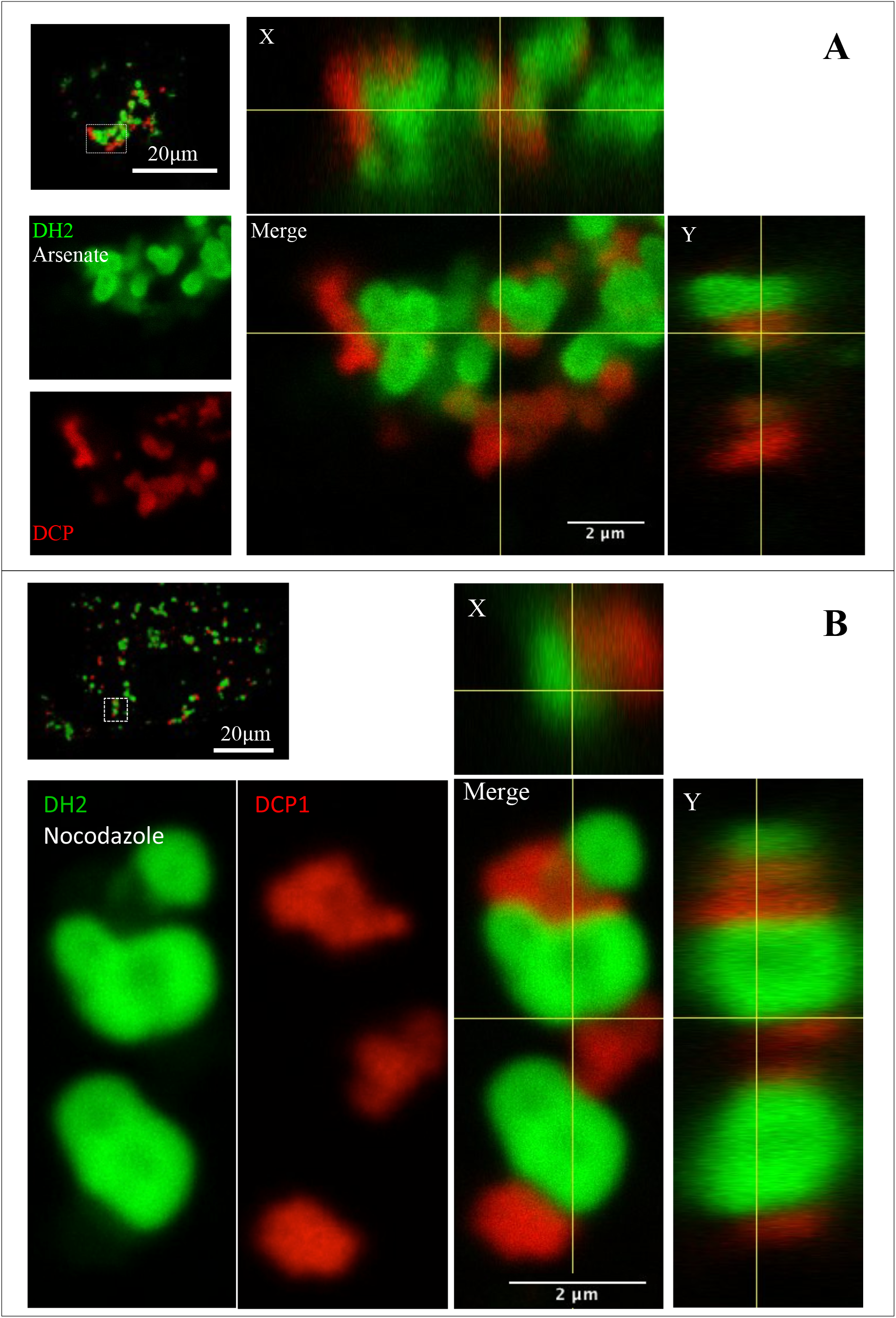
Cellular stress had no effect on formation of vesicle-like structures of DH2 and their ability to ‘trap’ DCP1. Confocal analysis of COS7 cells expressing GFP-tagged DH2 and mCherry-tagged DCP1 after arsenate (A) or nocodazole (B) treatment as described before. Merged images including vertical and side views (X and Y coordinates) shown on the right indicated large vesicle-like structures of DH2 adjacent to DCP1.

**Figure 7.**
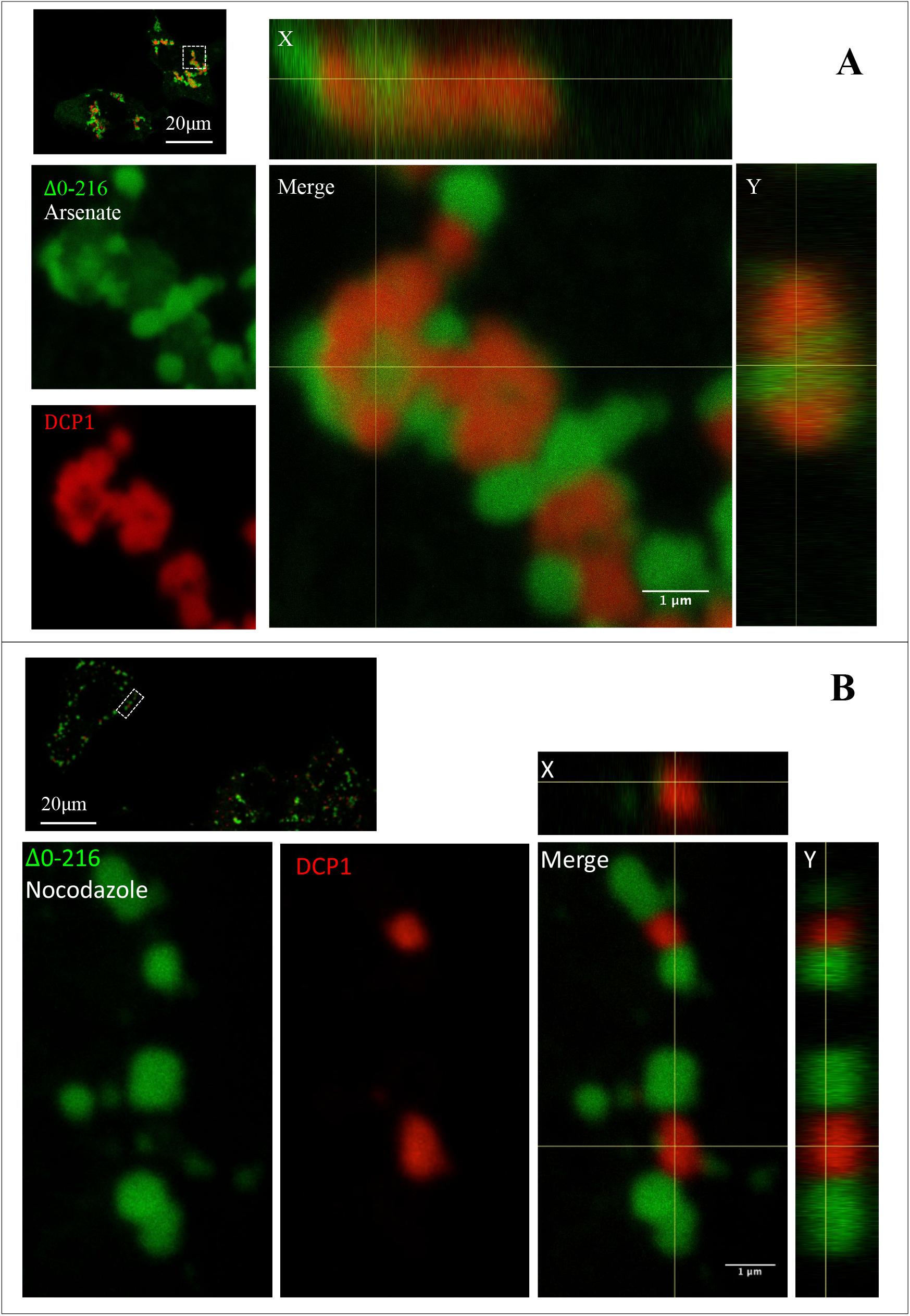
Cellular stress had no effect on formation of large structures of Δ0-216 and their ability to ‘trap’ DCP1. Confocal analysis of COS7 cells expressing GFP-tagged Δ0-216 and mCherry-tagged DCP1 after arsenate (A) or nocodazole (B) treatment as described before. Merged images including vertical and side views (X and Y coordinates) shown on the right indicated large structures of Δ0-216 adjacent to DCP1.

### ITAM and coiled coil domains are both required for colocalisation with P-bodies under cellular stress conditions

To further dissect the functional properties of the ITAM and the coiled coil domains, cellular localisation of a mutant protein just containing the ITAM domain or ITAM extent to PH domain (104-312aa), and the coiled coil disrupted mutants All-10 and Δ0-104-All-10 were tested after cellular stress. As shown in Figure 9, neither arsenate nor nocodazol treatment resulted in aggregates; rather all mutant proteins remained diffuse in the cytoplasm as in untreated cells similar to the GFP control. As shown before, wild type on the other hand did form granules colocalising with DCP1 after nocodazol treatment (Fig. 5). Together these results suggest that neither domain alone (ITAM or coiled coil) respond to cellular stress but in combination they do as shown above resulting in colocalisation with DCP1.

### Introduction of Y210/222E or Y210/222F mutations into Δ0-104 did not alter the morphology of aggregates but prevented the mutant proteins to completely colocalise with P-bodies under cellular stress condition

In the context of the full-length wild type DEF6 protein introduction of the phosphomimic Y210/222E mutations resulted in spontaneous granule formation and colocalisation with DCP1 (9) due to its unmasking effect on the N-terminal domain as shown in Cheng et al. (26). It was therefore not surprising that introduction of the same Y210/222E phosphomimic mutations into the Δ0-104 mutant lacking the N-terminal 104 amino acids did have no effect on the formation of aggregates that partially overlapped with DCP1 in untreated cells (Fig. 10). Similarly, the equivalent Y210/222F mutations introduced into Δ0-104 that prevent potential phosphorylation on Tyr 210 and 222 also did not show an effect (Suppl Fig. 1). However, in contrast to Δ0-104, introduction of either Y210/222E or Y210/222F into Δ0-104 prevented complete colocalisation with DCP1 after nocodazol treatment (Fig. 12). These results suggest that alteration in the ITAM (Y210E/F) and PH domain (Y222E/F) influence the structural conformation independent of the phosphorylation status that alters the response of Δ0-104 to cellular stress.

### The N- and C-terminal of DEF6 mediate interactions

As described above and in Cheng et al. (26), in isolation, both the N-terminal end as well as the coiled coil domain of DEF6 forms spontaneously either granules or aggregates indicating that both N- and C-terminal parts of DEF6 are capable of oligomerisation. Full-length wild type DEF6 protein on the other hand is mainly diffuse in the cytoplasm when overexpressed in COS7 cells. However, endogenous DEF6 in Jurkat T cells does form granules that overlap with DCP1 (30) suggesting that also the wild type DEF6 protein can oligomerise. To test this directly, a mCherry-tagged wild type DEF6 protein was also tagged with a SV40 nuclear localisation signal (NLS) (mCherry-NLS-DEF6; Suppl. Fig. 17). Although DEF6 contains a functional NLS (amino acid positions 330-341), as verified by the fact the PH2 mutant including the NLS is located exclusively in the nucleus (Fig. 2), the NLS is masked in the full length DEF6 protein which is largely absent from the nucleus (Fig. 1). mCherry-NLS-DEF6 however is exclusively found in the nucleus (Fig. 12 upper panel). If full length wild type DEF6 dimerises or oligomersies then cotransfection of GFP-DEF6 with mCherry-NLS-DEF6 could result in the retention of mCherry-NLS-DEF6 in the cytoplasm or the import of GFP-DEF6 into the nucleus. As shown Fig. 12 (lower panel), cotransfection of GFP-DEF6 with mCherry-NLS-DEF6 resulted in a clearly visible localisation of GFP-DEF6 in the nucleus completely overlapping with mCherry-NLS-DEF6. This is the first direct evidence that full length wild type DEF6 dimerises and/or oligomerises. To test whether the observed interaction is mediated through the coiled coil domain, mCherry-NLS-DEF6 was cotransfected with mutant proteins N-590, DH1 and DH2. Similar to full length wild type DEF6, mCherry-NLS-DEF6 dragged some of the mutant N-590, DH1 and in part DH2 proteins into the nucleus (Fig. 12). However, DH2 aggregates in the cytoplasm trapped some mCherry-NLS-DEF6 protein preventing it from entering the nucleus (Fig. 12). Confocal analysis revealed that mCherry-NLS-DEF6 retained in the cytoplasm was within the vesicle-like tubes formed by DH2 aggregates (Fig. 13). Furthermore, mutant proteins Q371P-A374P, L442P-E445P, All-10 all disrupting the coiled coil domain, as well as DHL-N which contains only a part of the coiled coil domain did not interact with mCherry-NLS-DEF6 (Fig. 14). Together these data suggest that oligomerisiation of DEF6 is indeed mediated through the coiled coil domain. However, when mCherry-NLS-DEF6 was cotransfected with N-108 lacking the coiled coil domain, mCherry-NLS-DEF6 was also partially retained in the cytoplasm colocalising with N-108 granules (Fig. 13). Importantly, phosphomimic mutant Y210/222E that spontaneously colocalises with DCP1 also retained mCherry-NLS-DEF6 in the cytoplasm (Fig. 12) suggesting that DEF6 is capable of dimer formation and/or oligomerisation in P-bodies. A summary of the data is collated in Table 1.

**Table 1.**
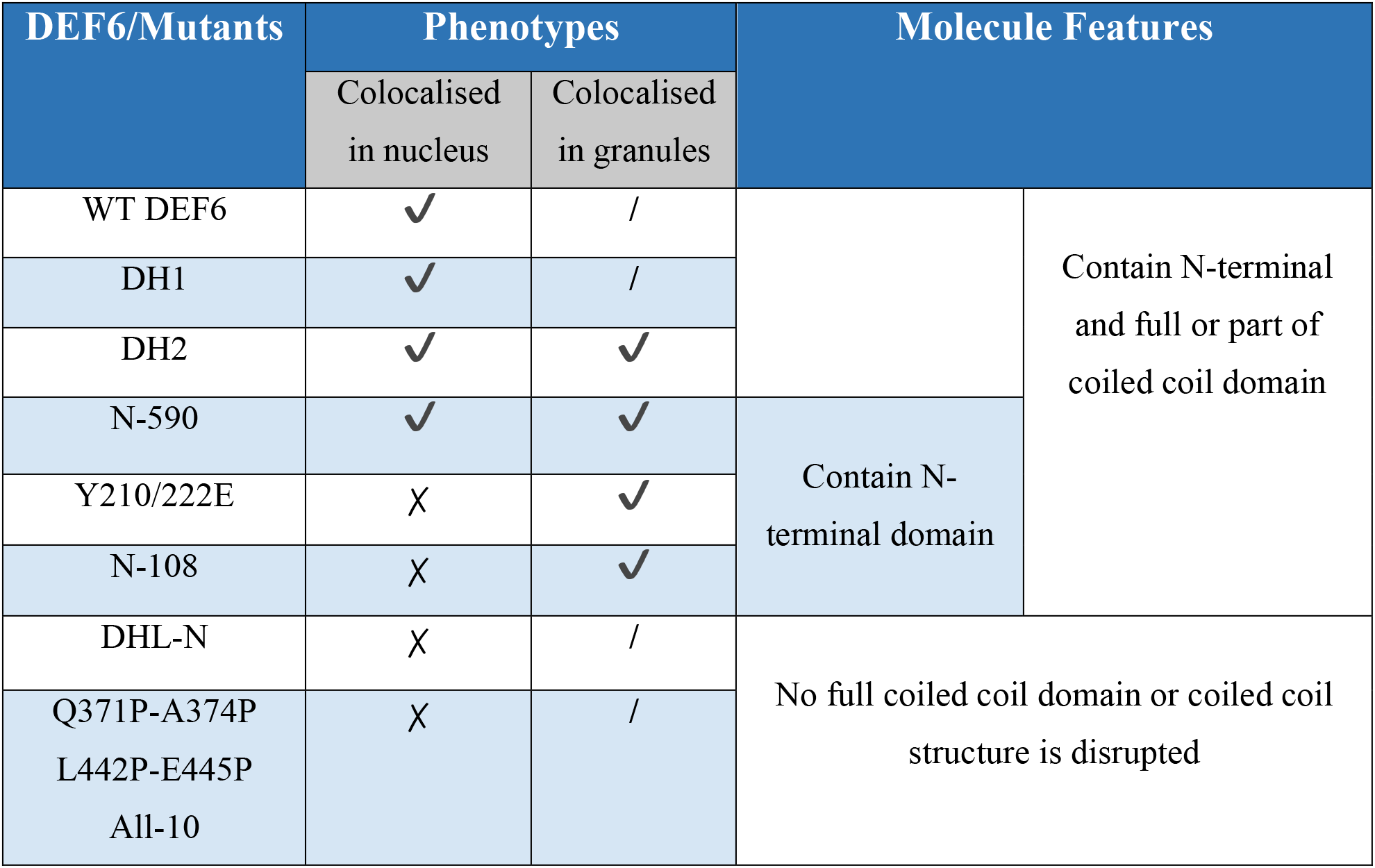
Cellular localisation of wild type and mutant DEF6 proteins coexpressed with mCherry-NLS-DEF6 Import of wild type or mutant proteins into the nucleus as well as retention of mCherry-NLS-DEF6 in the cytoplasm indicates interaction through dimerization and or oligomerisation. ✔:Yes, ✗:No, /: Not done

| DEF6/Mutants | Phenotypes |  | Molecule Features |  |
| --- | --- | --- | --- | --- |
|  | Colocalised in nucleus | Colocalised in granules |  |  |
| WT DEF6 | ✓ | / |  | Contain N-terminal and full or part of coiled coil domain |
| DH1 | ✓ | / |  |  |
| DH2 | ✓ | ✓ |  |  |
| N-590 | ✓ | ✓ | Contain N-terminal domain |  |
| Y210/222E | ✗ | ✓ |  |  |
| N-108 | ✗ | ✓ |  |  |
| DHL-N | ✗ | / | No full coiled coil domain or coiled coil structure is disrupted |  |
| Q371P-A374P<br>L442P-E445P<br>All-10 | ✗ | / |  |  |

## DISCUSSION

PairCoil2 analysis predicted a coiled coil domain in the glutamine (Q)-rich C-terminal part of DEF6 (between amino acids 330 and 550) (8). Q or hydrophobic amino acids occur frequently at the “a” and “d” position of the heptad repeats (a–g) of the predicted coiled coil region. Coiled coil domains can interact with other coiled coil domains resulting in right-handed supercoil structures that contain two to seven coiled coil domains (21). Given that wild type DEF6 tagged with GFP and overex pressed in COS7 cells localised diffuse in the cytoplasm, it is likely that the conformation of DEF6 is such that the coiled coil domain is masked to control its ability to interact with other coiled coil domains. However, several mutant versions of DEF6 (e.g. DH2, Y133/144D, Δ0-104, Δ0-216) formed aggregates in COS7 cells suggesting a conformational change unmasking the coiled coil domain. Some aggregates formed large structures that in the extreme were vesicle-like (mutant DH2; Fig. 4). It seems therefore that the coiled coil domain of DEF6 can interact with itself and if uncontrolled, this interaction results in large aggregates. Cotransfection of mutant Y133/144D with either DH2, Δ0-104 or Δ0-216 confirmed this interpretation: aggregates formed were completely overlapping indicating that DEF6 molecules can interact with each other via the coiled coil domain (Suppl. Fig. 2). While the N-terminal 45 amino acids are sufficient, the first 104 amino acids are not required for P-body colocalisation. Δ0-104 lacking the first 104 amino acids but containing ITAM, PH and the coiled coil domain (Fig. 1A), formed coiled coil-mediated aggregates that trapped DCP1 in untreated COS7 cells, but formed granules that completely overlapped with DCP1 in arsenate or nocodazole treated cells (Fig. 8) indicating that stress induced modifications of Δ0-104 results in conformational change unmasking parts of DEF6 within ITAM, PH and/or coiled coil domains that facilitates P-body colocalisation. However, neither domain on its own colocalised with P-bodies. ITAM and PH1 localised diffuse in the cytoplasm and nucleus and DH2 containing the coiled coil domain formed aggregates (Fig. 2) suggesting that the stress-induced conformation of Δ0-104 is required for P-body localisation. Indeed, introduction of Y210/222E into Δ0-104 did not change aggregate formation in untreated cells but prevented complete overlap with DCP1 in stressed cells (Fig. 11). This might suggest that ITK phosphomimic mutant in the context of the N-terminal truncation mutant Δ0-104 ‘fixed’ its conformation preventing stress-induced modifications. It also implies the phosphorylation on DEF6 Y210/222 is more likely respond to the N-terminal mediated P-body localisation rather than the C-terminal (26). Overall, these results suggest that ITAM, PH and coiled coil can cooperate in a stress-induced conformation that facilitates P-body localisation independent of the N-terminal end. The experiments described above also showed that mutant DEF6 proteins forming coiled coil-mediated aggregates (DH2; Y210/222E) can alter the behaviour of wild type DEF6 that now also formed aggregates colocalising with the mutant protein (Fig. 12). Either the coiled coil domains of the mutant proteins in the aggregates are able to interact with the coiled coil domain of wild type DEF6 trapping it in the aggregates or the mutant protein enforced a conformational change that resulted in aggregation of the wild type protein. The latter interpretation would mean that mutant DEF6 can act prion-like, a notion that is supported by the finding that Y133/144D mutant that spontaneously aggregates can trigger aggregations of other proteins (537-590 and 537-550) that on their own do not aggregate (31). However, some aggregates formed by mutant proteins trapped mCherry-tagged DCP1 that no longer exhibited its normal cytoplasmic distribution when cotransfected (e.g. DH2 and Δ0-216; Fig. 4; Y133/144D; Suppl. Fig.1). DCP1 does not contain a coiled coil domain but many P-body components do and it remains to be seen whether DCP1 on its own was trapped by the aggregates or whether localisation of entire P-bodies was altered.

**Figure 8.**
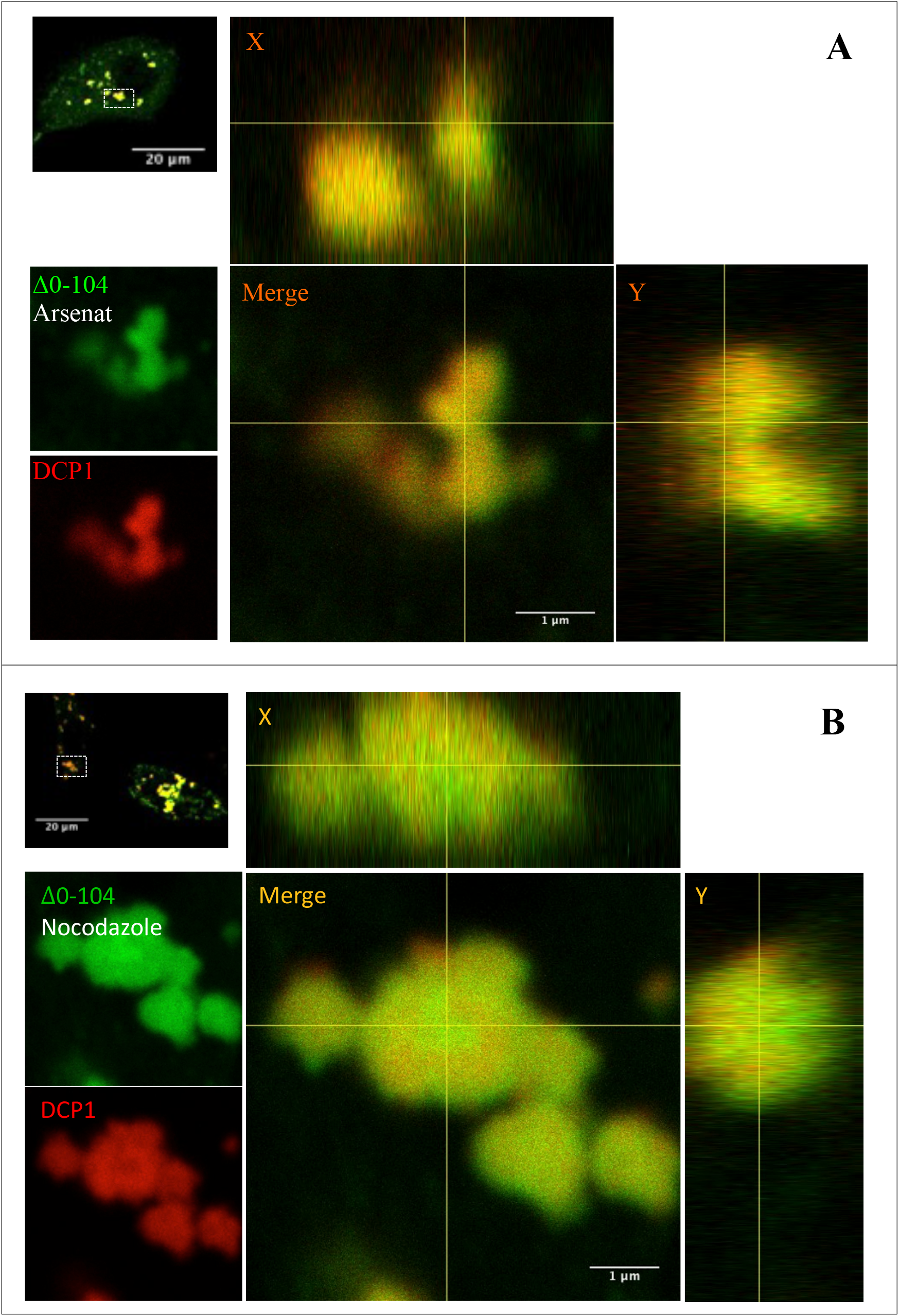
Cellular stress results in complete colocalisation of Δ0-104 structures with DCP1. Confocal analysis of COS7 cells expressing GFP-tagged Δ0-104 (representing group 3 mutants) and mCherry-tagged DCP1 after arsenate (A) or nocodazole (B) treatment as described before. Merged images including vertical and side views (X and Y coordinates) shown on the right indicated that large structures of Δ0-104 completely overlapped with DCP1.

**Figure 9.**
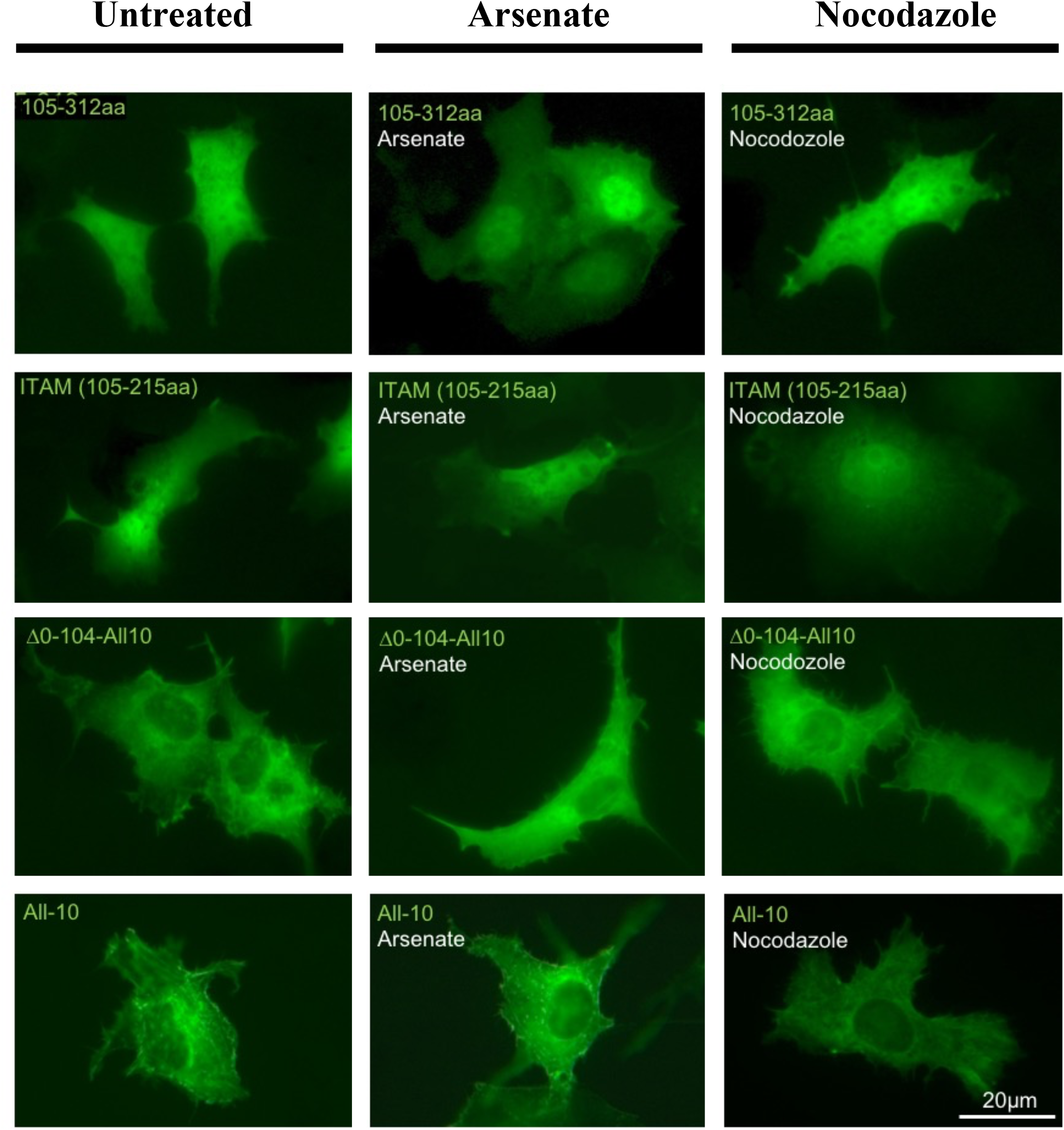
Cellular stress did not alter localisation of the mutant protein 105-312aa, ITAM (105-215aa), Δ0-104-A11-10 and All-10. Comparison of nontreated (left panel) with either arsenate (middle panel) or nocodazole (right panel) treated cells transfected with the DEF6 proteins as indicated revealed that a functional coiled coil domain is required for the formation of aggregates even under cellular stress conditions which in contrast to mutants of the 3^rd^ group, e.g. Δ0-104 (see Fig. 1,3 and 4).

**Figure 10.**
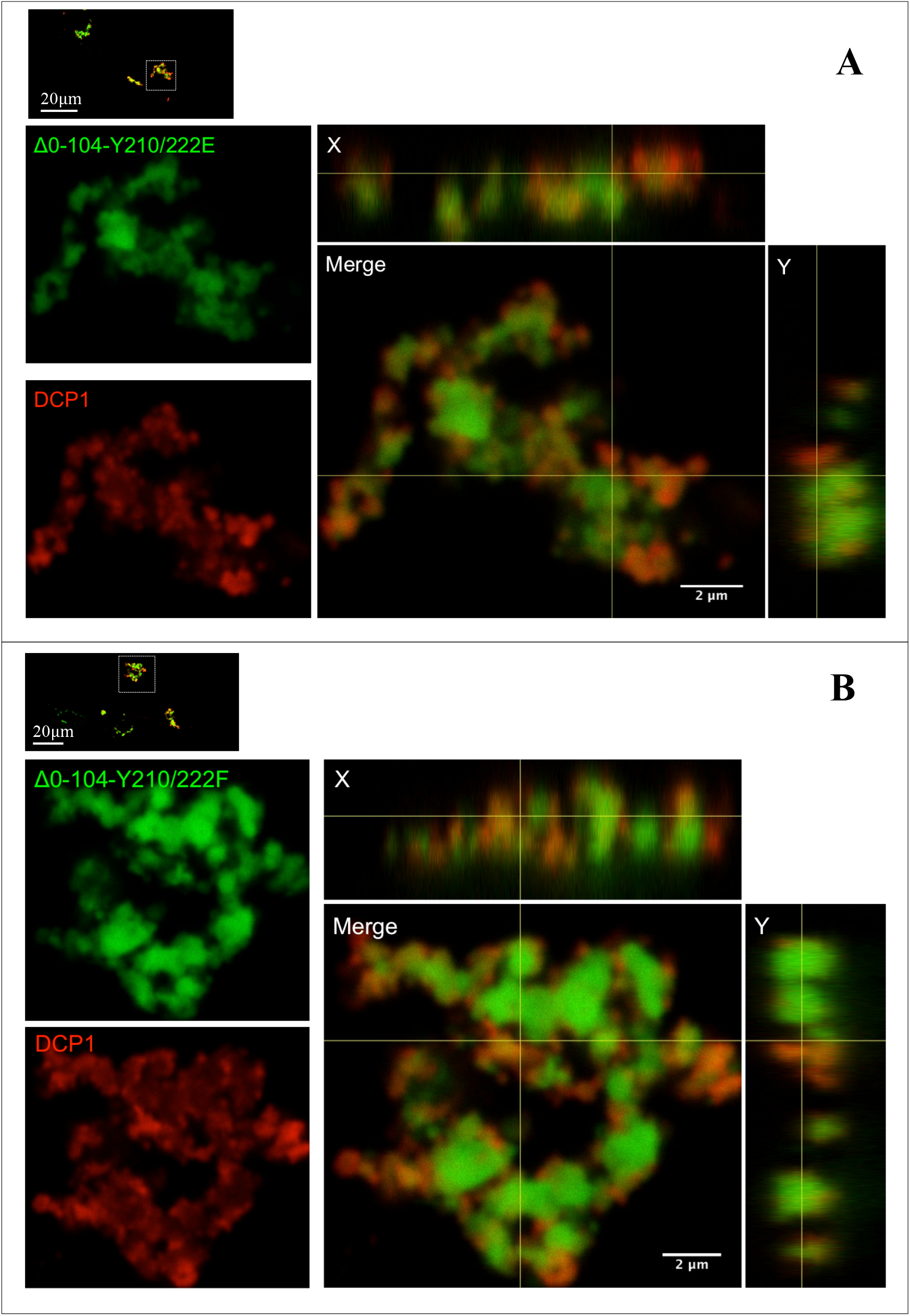
Introduction of Y210/222E or Y210/222F mutations into Δ0-104 did not alter the morphology of aggregates that still partially overlapped with DCP1. Confocal analysis of COS7 cells expressing GFP-tagged Δ0-104-Y210/222E (A) or Δ0-104-Y210/222F (B) and mCherry-tagged DCP1 as described before. Merged images including vertical and side views (X and Y coordinates) shown on the right indicated that both mutant proteins formed large structures partially overlapped with DCP1.

**Figure 11.**
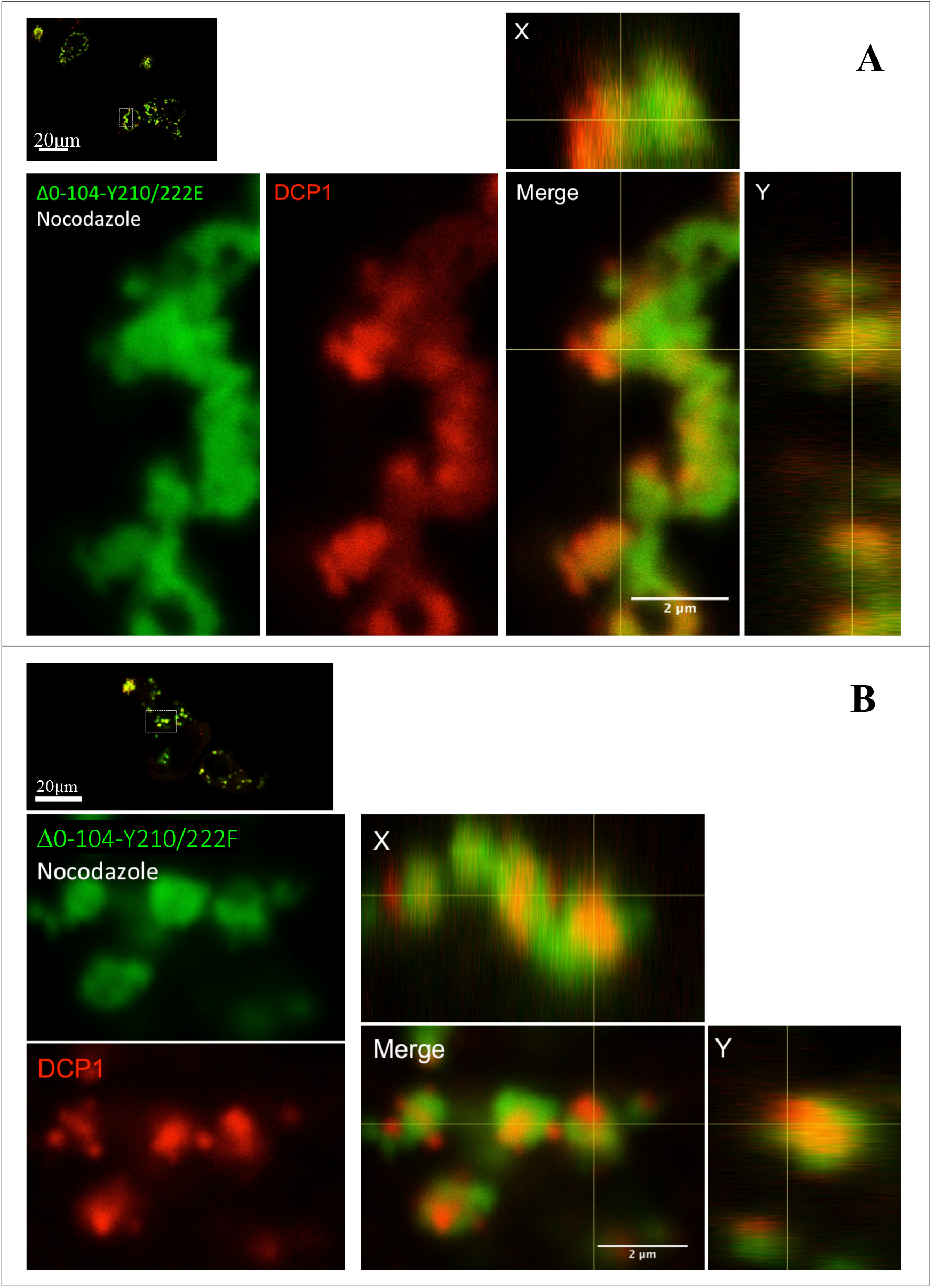
Introduction of Y210/222E or Y210/222F mutations into Δ0-104 prevented complete overlap with DCP1 under cellular stress conditions. Confocal analysis of COS7 cells expressing GFP-tagged Δ0-104-Y210/222E (A) or Δ0-104-Y210/222F (B) and mCherry-tagged DCP1 after nocodazole treatment as described before. Merged images including vertical and side views (X and Y coordinates) shown on the right indicated that both mutant proteins formed large structures partially overlapped with DCP1 which differs from Δ0-104 that completely overlapped with DCP1 under stress conditions (see Figure 8).

**Figure 12.**
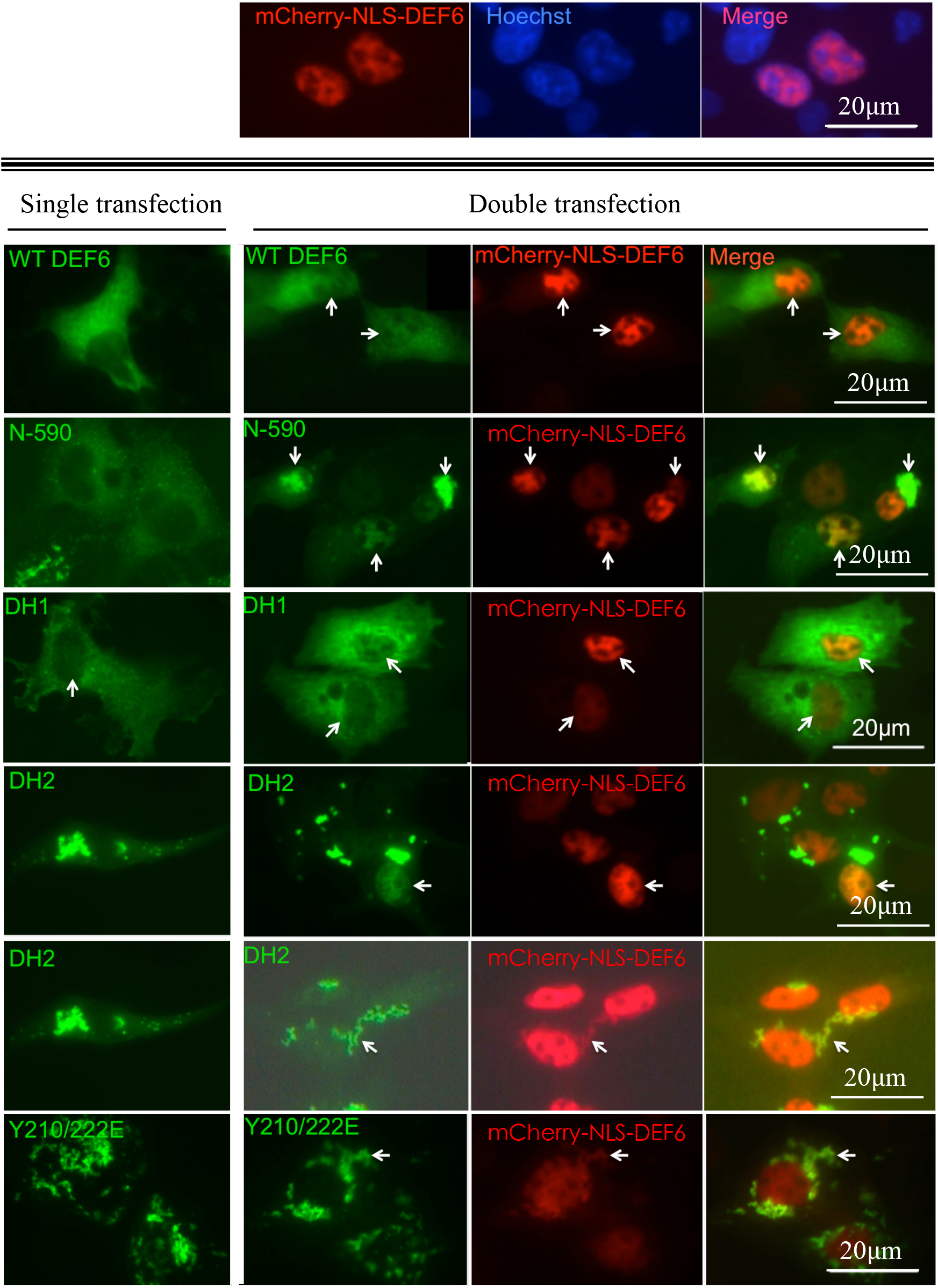
Interaction of DEF6 molecules through dimerisation and or oligomerisation. Upper Panel: mCherry-NLS-DEF6 localised exclusively to the nucleus that was stained with Hoechst. Lower Panel: WT DEF6, N-590, DH1 and DH2 that on their own were localised in the cytoplasm (left column) as shown before, cotransfection with mCherry-NLS-DEF6 resulted in clear nuclear import of these mutants (merged images in the right column). In addition, DH2 and Y210/222E aggregations (lower two rows) retained some mCherry-NLS-DEF6 in the cytoplasm.

**Figure 13.**
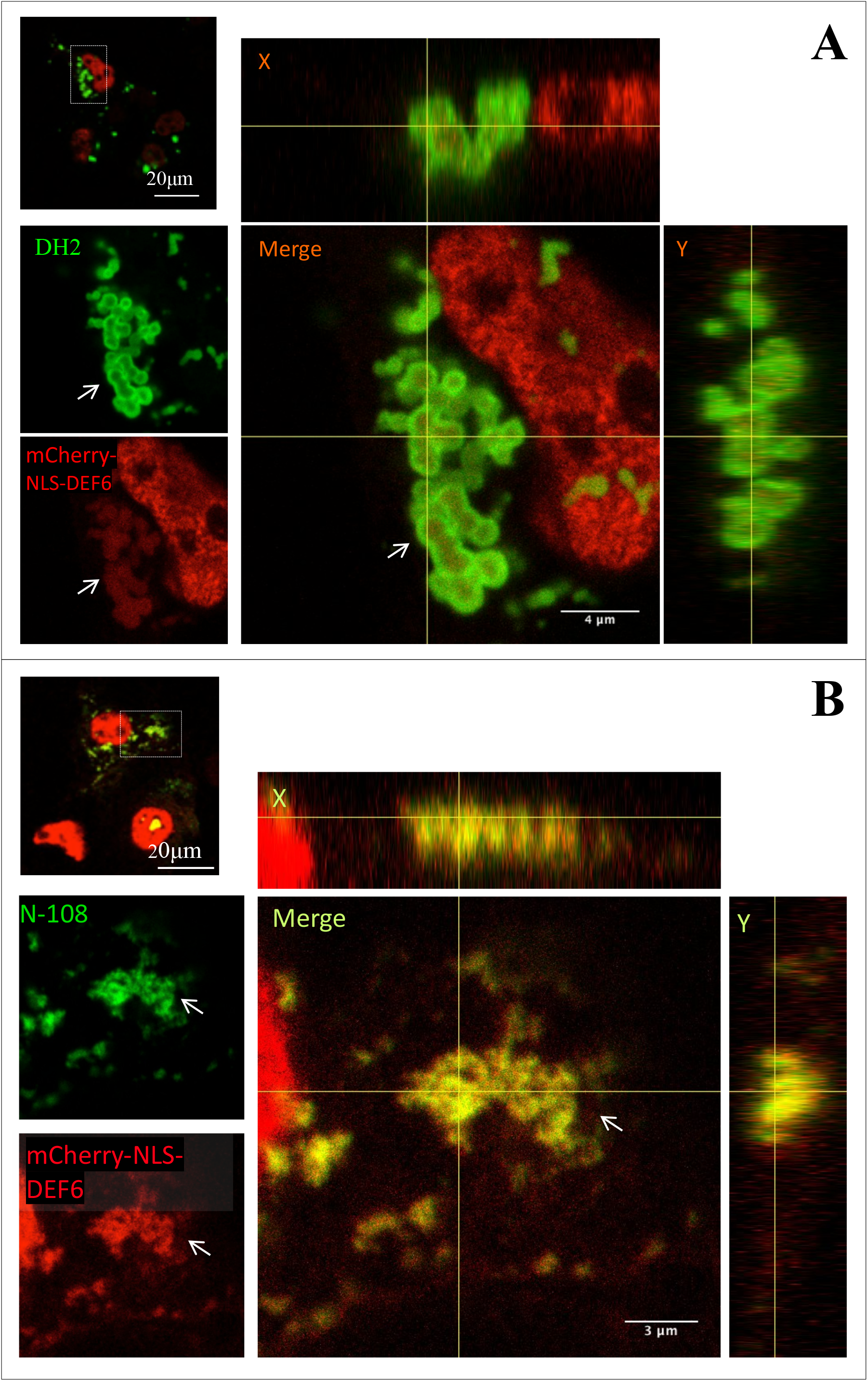
Dimerisation and or oligomerisation of DEF6 can also be mediated through its N-terminal 108 amino acids. Confocal analysis of COS7 cells cotransfected with mCherry-NLS-DEF6 and either DH2 (A) or N-108 (B) as described before. Merged images on the right including vertical and side views (X and Y coordinates) show that DH2 vesicle-like structures retain some mCherry-NLS-DEF6 in the cytoplasm by ‘trapping’ it within the vesicle-like structures. N-108 also formed cytoplasmic aggregations that retained some mCherry-NLS-DEF6 in the cytoplasm indicating the the N-terminal 108 amino acids of DEF6 can also facilitate interaction resulting demerisation and or oligomerisation.

**Figure 14.**
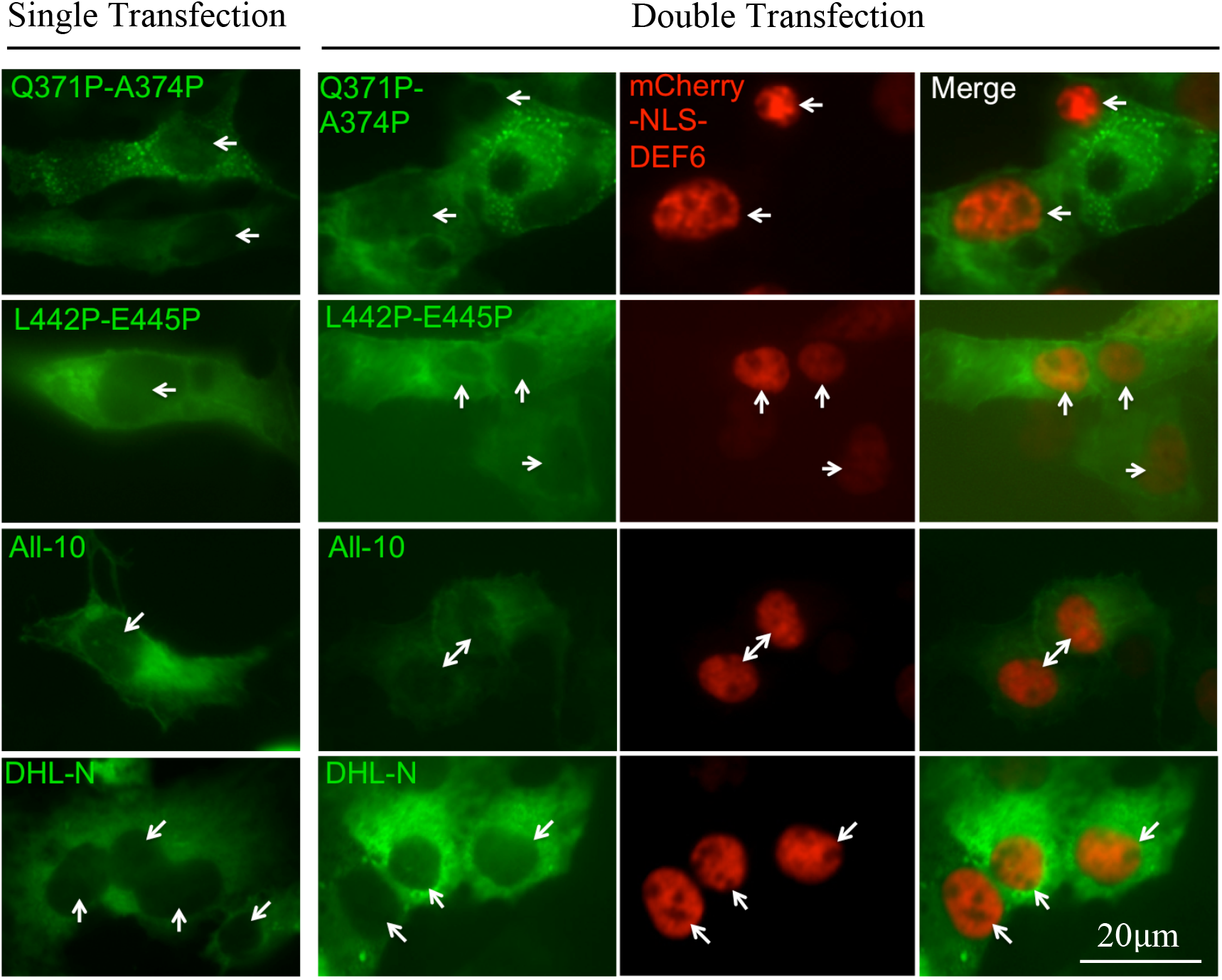
DEF6 mutants Q371P-A374P, L442P-E445P, All-10 and DHL-N did not interact with mCherry-NLS-DEF6. Comparison of single transfection (left column) and cotransfection of Q371P-A374P, L442P-E445P, All-10 and DHL-N mutants with mCherry-NLS-DEF6 (right three columns). Merged images on the right show that mutants with disrupted coiled coil and/or partial coiled coil domain failed to interact with mCherry-NLS-DEF6.

## EXPERIMENTAL PROCEDURES

WT and mutant DEF6 tagged with GFP or mCherry were expressed in COS7 cells and their cellular localization analysed as describe (26). mCherry-SiT-N-15 plasmid was a gift from Michael Davidson (Addgene plasmid # 55133; http://n2t.net/addgene:55133;RRID:Addgene_55133). mCherry tagged GalT was generated from Cerulean-GalT plasmid by replacing cerulean to mCherry using subcloning technology, and the Cerulean-GalT plasmid was a gift from Jennifer Lippincott-Schwartz (Addgene plasmid # 11930; http://n2t.net/addgene:11930;RRID:Addgene_11930). mCherry-NLS-DEF6 plasmid was establish by inserting annealed NLS oligos (ThermoFisher Scientific) between mCherry and DEF6, and mCherry tagged DEF6 was generated by subcloning technology. All the mutants were verified by sequencing (Suppl. Fig. 14 to 18). The Nocodazole (Sigma, #M1404-2MG) treatment was carried out by 1 *μ*m/ml final concentration, and the cells were treated in their common culture medium in 37° C for 2 hours.

## Acknowledgments

We thank Drs. Martin Gering, Sally Wheatley and Peter Jones for helpful discussions, the SLIM team at the University of Nottingham for their support in confocal microscopy and the undergraduate students George Harrison Church, Sarah Mcginley and Chris Young for their assistants in establishing some of the DEF6 mutants.

## Conflict of interest

The authors declare that they have no conflicts of interest with the contents of this article.JBC Editorial Policies.

## FOOTNOTES

The abbreviations used are: DEF6, Differentially Expressed in FDCP 6; IBP, IRF4-binding protein; SLAT, SWAP-70-like adaptor protein of T cells; IS, immunological synapse; P-body, mRNA processing body; DCP1, decaping protein 1, ITAM, immunoreceptor tyrosine-based activation motif like sequence; PH domain, pleckstrin-homology domain; DHL domain, Dbl homology-like domain; GEF, guanine nucleotide exchange factor.

## REFERENCE

1. Hotfilder, M., Baxendale, S., Cross, M.A., Fred. S. (1999). Def-2, −3, −6 and −8, novel mouse genes differentially expressed in the haemopoietic system. British Journal of Haematology, 106, 335–344.

2. Borggrefe, T., Wabl, M., Akhmedov, A. T., & Jessberger, R. (1998). A B-cell-specific DNA recombination complex. Journal of Biological Chemistry, 273(27), 17025–17035.

3. Gupta, S., Lee, A., Hu, C., Fanzo, J., Goldberg, I., Cattoretti, G., & Pernis, A. B. Molecular cloning of IBP, a SWAP-70 homologous GEF, which is highly expressed in the immune system. Human Immunology, 64(4), 389–401.

4. Ripich, T., Chacón-Martínez, C. A., Fischer, L., Pernis, A., Kiessling, N., Garbe, A. I., & Jessberger, R. (2016). SWEF proteins distinctly control maintenance and differentiation of hematopoietic stem cells. PLoS ONE, 11(8), 1–12.

5. Manni, M., Ricker, E., & Pernis, A. B. (2017). Regulation of systemic autoimmunity and CD11c + Tbet + B cells by SWEF proteins. Cellular Immunology, (April), 0–1.

6. Shuen, W. H. (2010) Molecular Evolution of the def6/swap70 Gene Family and Functional Analysis of swap70a in Zebrafish Embryogenesis, MRes Thesis, The University of Nottingham

7. Mavrakis, K. J., McKinlay, K. J., Jones, P., & Sablitzky, F. (2004). DEF6, a novel PH-DH-like domain protein, is an upstream activator of the Rho GTPases Rac1, Cdc42, and RhoA. Experimental Cell Research, 294(2), 335–44.

8. Bécart, S., Balancio, A. J. C., Charvet, C., Feau, S., Sedwick, C. E., & Altman, A. (2008). Tyrosine-phosphorylation-dependent translocation of the SLAT protein to the immunological synapse is required for NFAT transcription factor activation. Immunity, 29(5), 704–19.

9. Hey, F., Czyzewicz, N., Jones, P., & Sablitzky, F. (2012). DEF6, a novel substrate for the Tec kinase ITK, contains a glutamine-rich aggregation-prone region and forms cytoplasmic granules that co-localize with P-bodies. The Journal of Biological Chemistry, 287(37), 31073–84.

10. Gupta, S., Fanzo, J. C., Hu, C., Cox, D., Jang, S. Y., Lee, A. E., … Pernis, A. B. T cell receptor engagement leads to the recruitment of IBP, a novel guanine nucleotide exchange factor, to the immunological synapse. The Journal of Biological Chemistry, 278(44), 43541–9.

11. Jakymiw, A., Lian, S., Eystathioy, T., Li, S., Satoh, M., Hamel, J. C., … Chan, E. K. L. (2005). Disruption of GW bodies impairs mammalian RNA interference. Nature Cell Biology, 7(12), 1267–74.

12. Eulalio, A., Behm-Ansmant, I., & Izaurralde, E. (2007). P bodies: at the crossroads of post-transcriptional pathways. Nature Reviews Molecular Cell Biology, 8(1), 9–22.

13. Anderson, P., & Kedersha, N (a). (2008). Stress granules: the Tao of RNA triage. Trends in Biochemical Sciences, 33(3), 141–150.

14. Anderson, P., & Kedersha, N (b). (2009). Stress granules. Current Biology, 19(C), 397–398.

15. Chang, C. Te, Bercovich, N., Loh, B., Jonas, S., & Izaurralde, E. (2014). The activation of the decapping enzyme DCP2 by DCP1 occurs on the EDC4 scaffold and involves a conserved loop in DCP1. Nucleic Acids Research, 42(8), 5217–5233.

16. Parker, R., & Sheth, U. (2007). P Bodies and the Control of mRNA Translation and Degradation. Molecular Cell, 25(5), 635–646.

17. Wang, C.-Y., Chen, W.-L., & Wang, S.-W. (2013). Pdc1 Functions in the Assembly of P Bodies in *Schizosaccharomyces pombe*. Molecular and Cellular Biology, 33(6), 1244–1253.

18. Rybak, A., Fuchs, H., Hadian, K., Smirnova, L., Wulczyn, E. A., Michel, G., … Wulczyn, F. G. (2009). The let-7 target gene mouse lin-41 is a stem cell specific E3 ubiquitin ligase for the miRNA pathway protein Ago2. Nature Cell Biology, 11(12), 1411–1420.

19. Decker, C. J., & Parker, R. (2012). P-Bodies and Stress Granules: Possible Roles in the Control of Translation and mRNA Degradation. Cold Spring Harbor Perspectives in Biology, 1–16.

20. Ferdinando Fiumara, Luana Fioriti, Eric R. Kandel, W. A. H. (2012). Essential role of coiled-coils for aggregation and activity of Q/N- rich prions and polyQ proteins, Cells, 143(7), 1121–1135.

21. Liu, J., Zheng, Q., Deng, Y., Cheng, C., Kallenbach, N. R., & Lu, M. (2006). A seven-helix coiled coil. PNAS, 103(42).

22. Wolf, E., Kim, P. S., & Berger, B. (1997). Multicoil - a program for predicting 2-stranded and 3-stranded coiled coils. Protein Sci, 6(6), 1179–1189.

23. Mason, J. M., & Arndt, K. M. (2004). Coiled coil domains: Stability, specificity, and biological implications. ChemBioChem, 5(2), 170–176.

24. Mier, P., Alanis-Lobato, G., & Andrade-Navarro, M. A. (2017). Protein-protein interactions can be predicted using coiled coil co-evolution patterns. Journal of Theoretical Biology, 412(November 2016), 198–203.

25. Mollett, E. (2014) A study of DEF6 granule formation using biophysical and cellular methods, PhD Thesis, University of Nottingham

26. Cheng, H., Alsayegh, M., and Sablitzky, F (2019) N-terminal 45 amino acids of DEF6 are necessary and sucientto spontaneously colocalise with DCP1 in P-bodies. bioRxiv doi: http://dx.doi.org/10.1101/533109

27. Martin, P. (2007). Analysis of Def6, a novel guanine nucleotide exchange factor showing dynamic regulation in vitro and playing an essential role in zebrafish development, PhD Thesis, University of Nottingham.

28. Sweet, T. J., Boyer, B., Hu, W., Baker, K. E., & Coller, J. (2007). Microtubule disruption stimulates P-body formation. Rna, 13(4), 493–502.

29. Carbonaro, M., O’Brate, A., & Giannakakou, P. (2011). Microtubule disruption targets HIF-1*α* mRNA to cytoplasmic P-bodies for translational repression. Journal of Cell Biology, 192(1), 83–99.

30. Remon K.L. (2016) DEF6 Aggregation is Linked to Active Translation and mRNA Turnover in T Cells, PhD Thesis, University of Nottingham

31. Huaitao Cheng and Fred Sablitzky (in preparation)

