## Supplementary material for "The coiled coil domain of DEF6 facilitates formation of large vesicle-like, cytoplasmic aggregates that trap the P-body marker DCP1 and exhibit prion-like features": Suppl.

### LCK phosphomimic mutant Y133-144D forms aggregates that interact with DEF6 N-terminal truncated mutants

It had been reported that the LCK phosphorylation of DEF6 is required for DEF6 to locate to the IS and that phosphomimic mutant Y133/144D spontaneously localised to the IS (8, 10). However, expression of Y133/144D in COS7 cells resulted in aggregate formation that partially overlapped with DCP1 (Suppl. Fig. 1). This phenotype is reminiscent of the one observed with DEF6 N-terminal truncated mutants such as Δ0-104 (Fig. 4). However, unlike to Δ0-104 (Fig. 8), nocodazole treatment had no effect with Y133/144D still only partially overlapping with DCP1 (Suppl. Fig. 1). mCherry-tagged Y133/144D aggregates did however fully colocalise with aggregates formed by DH2, Δ0-216 and Δ0-104 (Suppl. Fig. 2).

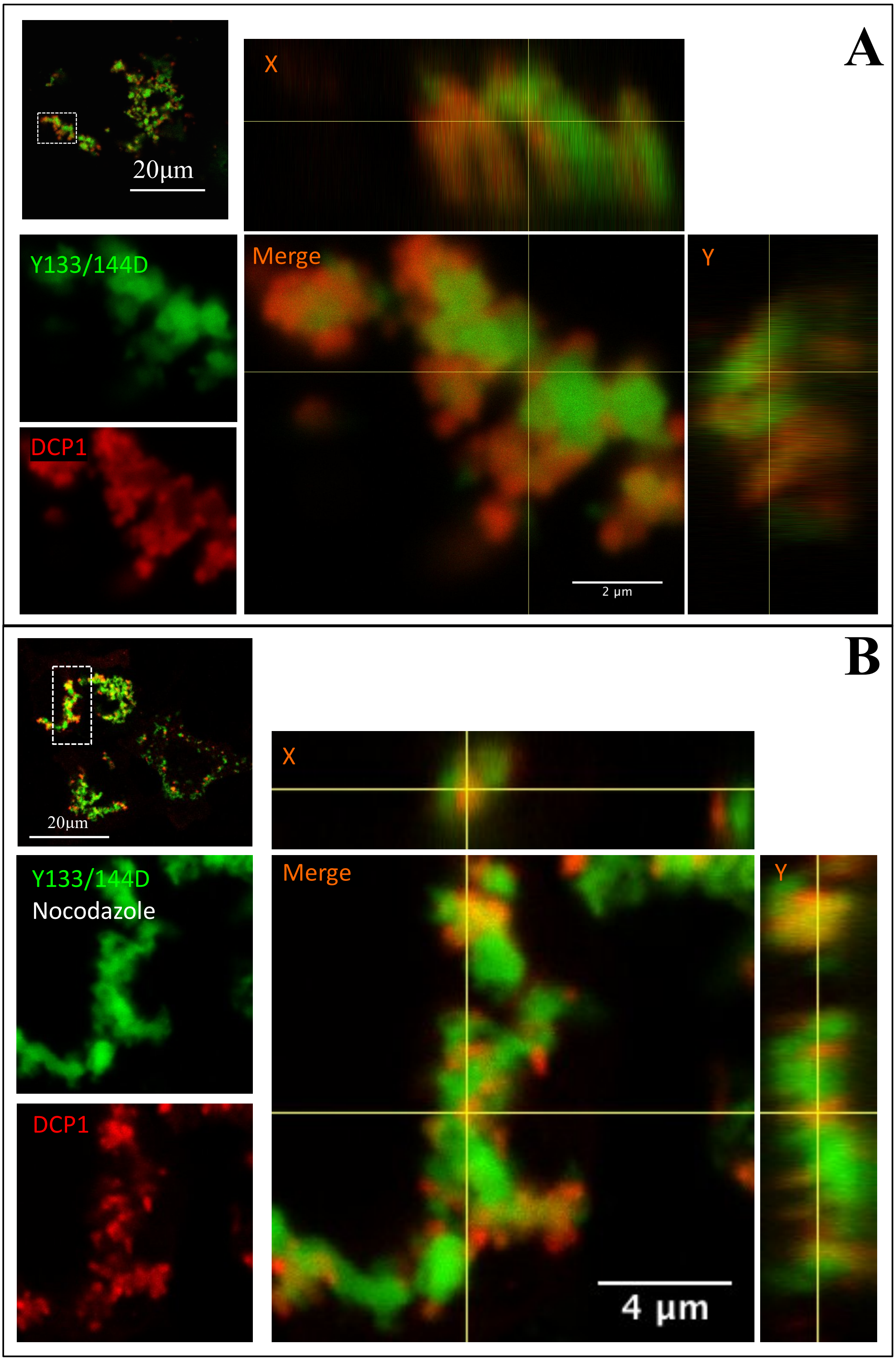
A
20µm
X
Y133/144D
Merge
Y
DCP1
2 µm
B
20µm
X
Y133/144D
Nocodazole
Merge
Y
DCP1
4 µm

**Supplementary Figure 1.** LCK phosphomimic mutant Y133/144D formed aggregates that partially overlapped with DCP1 in untreated and nocodazole treated cells. Confocal analysis of COS7 cells cotransfected with GFP-tagged Y133/144D and mCherry-tagged DCP1 in untreated cells (A) and after nocodazole treatment (B). Boxed area in the upper left images are enlarged in the lower left and further enlarged in the merged images on the right. Vertical and side views (X and Y coordinates) are also shown on the right. DCP1 is trapped by Y133/144D aggregates resulting in partial overlap in untreated and nocodazole treated cells.

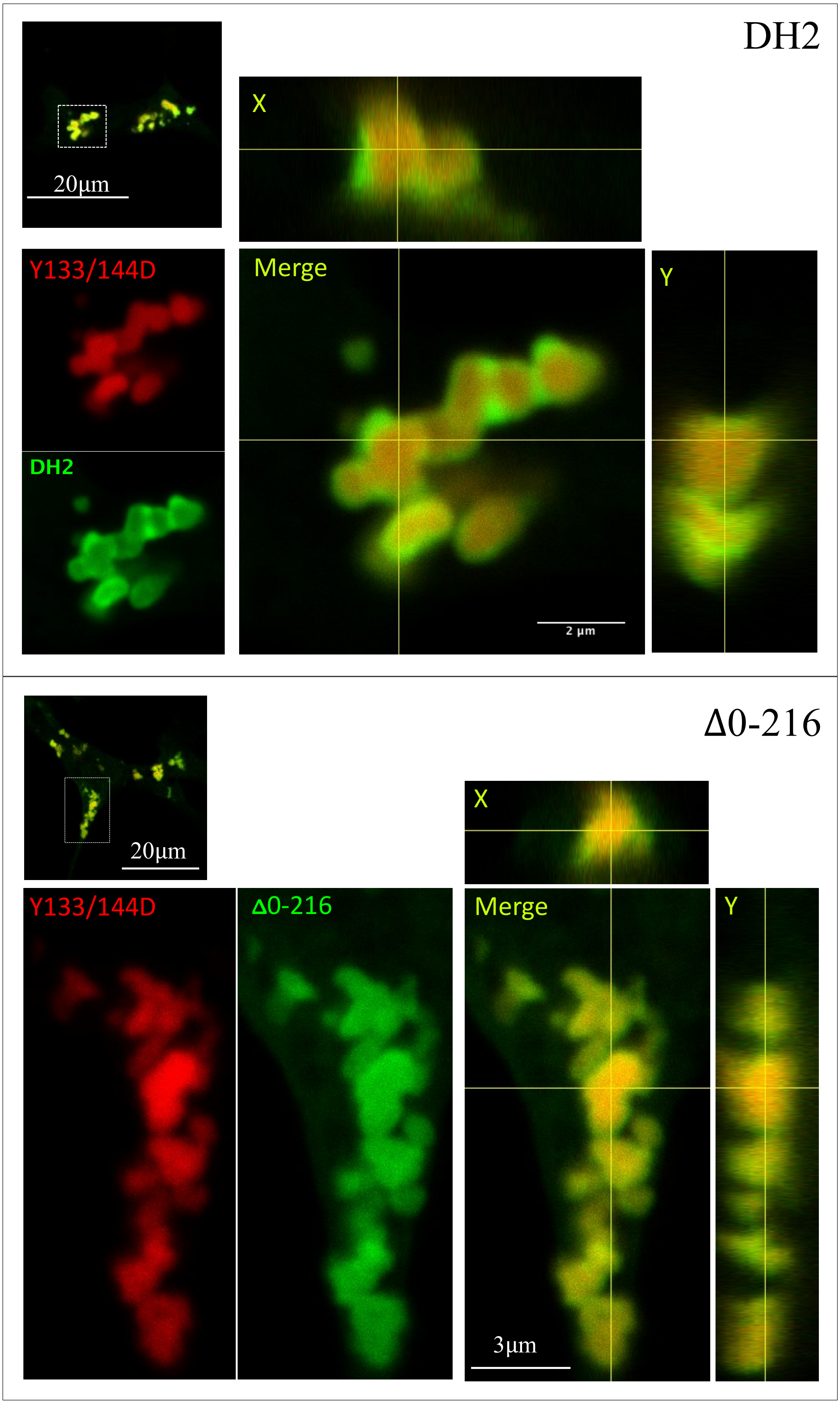
DH2
20μm
X
Y133/144D
Merge
Y
DH2
2 μm
Δ0-216
20μm
X
Y133/144D
Δ0-216
Merge
Y
3μm

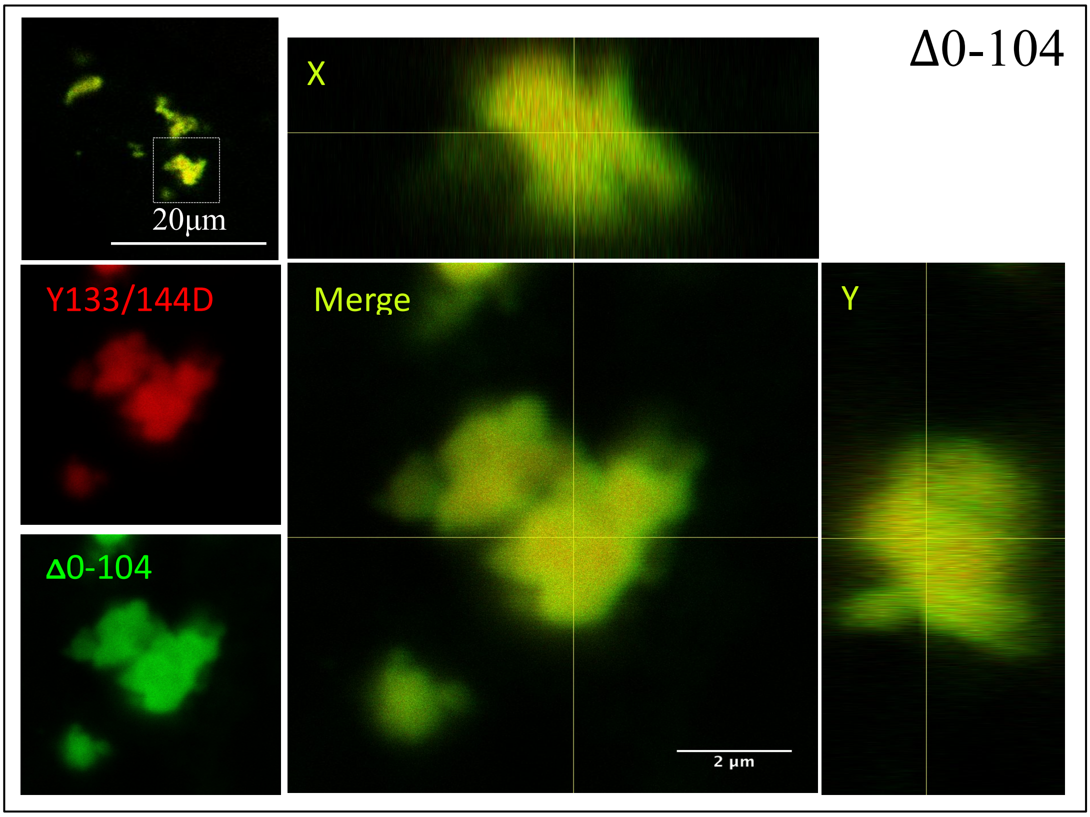


**Supplementary Figure 2.** Aggregates formed by the LCK phosphomimic mutant Y133/144D interact with DH2, Δ0-216 and Δ0-104. Confocal analysis of COS7 cells cotransfected with mCherry-tagged Y133/144D (red) and either GFP-tagged DH2 (upper panel), Δ0-216 (middle panel) or Δ0-104 (lower panel; green). Boxed area in the upper left images are enlarged in the lower left and further enlarged in the merged images on the right. Vertical and side views (X and Y coordinates) are also shown on the right in each panel. Complete overlap of aggregates formed indicate that Y133/144D mutant aggregates are mediated through the coiled coil domain.

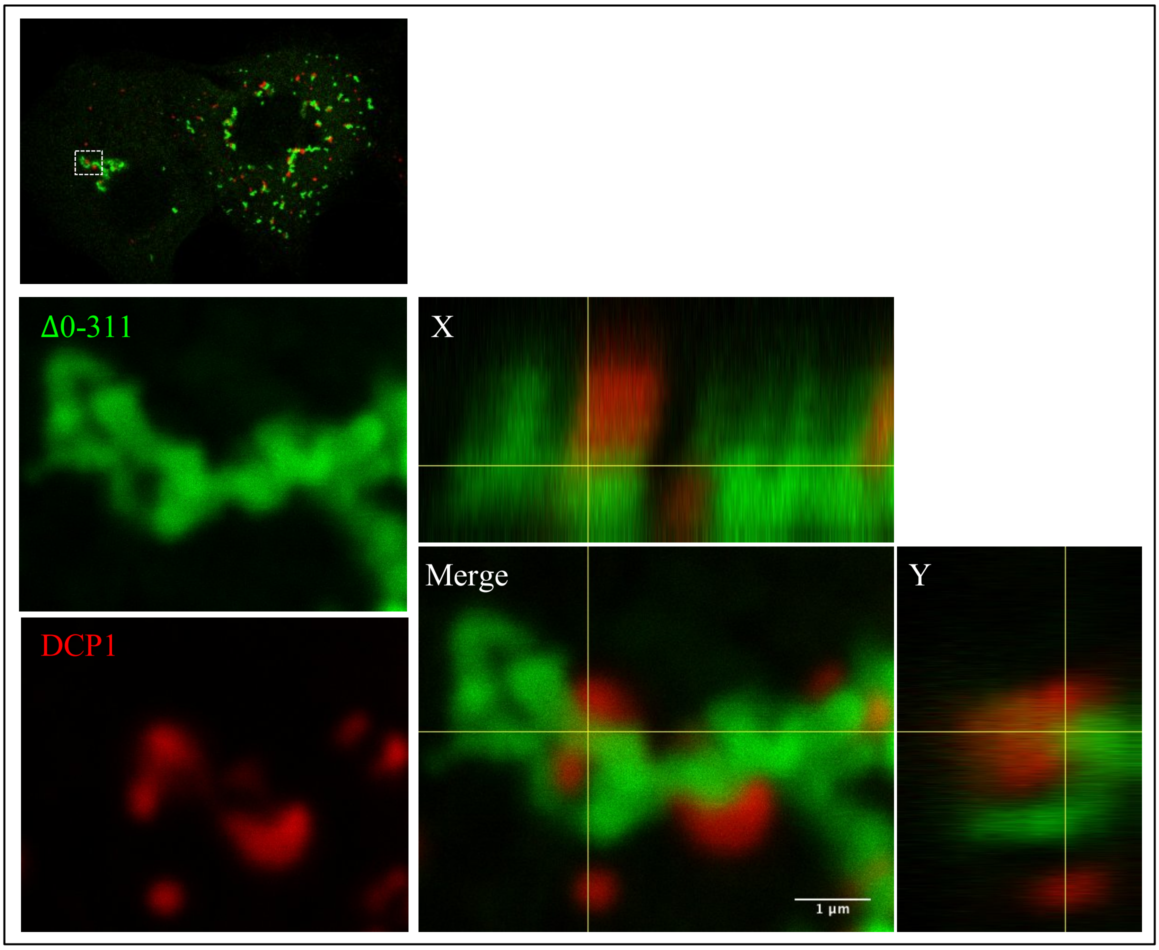


**Supplementary Figure 3.** Coiled coil-mediated aggregates (Δ0-311) of DEF6 form large structure that 'trap' DCP1. Confocal analysis of transfected COS7 cells as described before. GFP-tagged Δ0-311 aggregates also formed large structures but these did not appear vesicle-like. However, these structures also altered the localisation of DCP1 that was always associated with DEF6 structures (see merged images on the right)

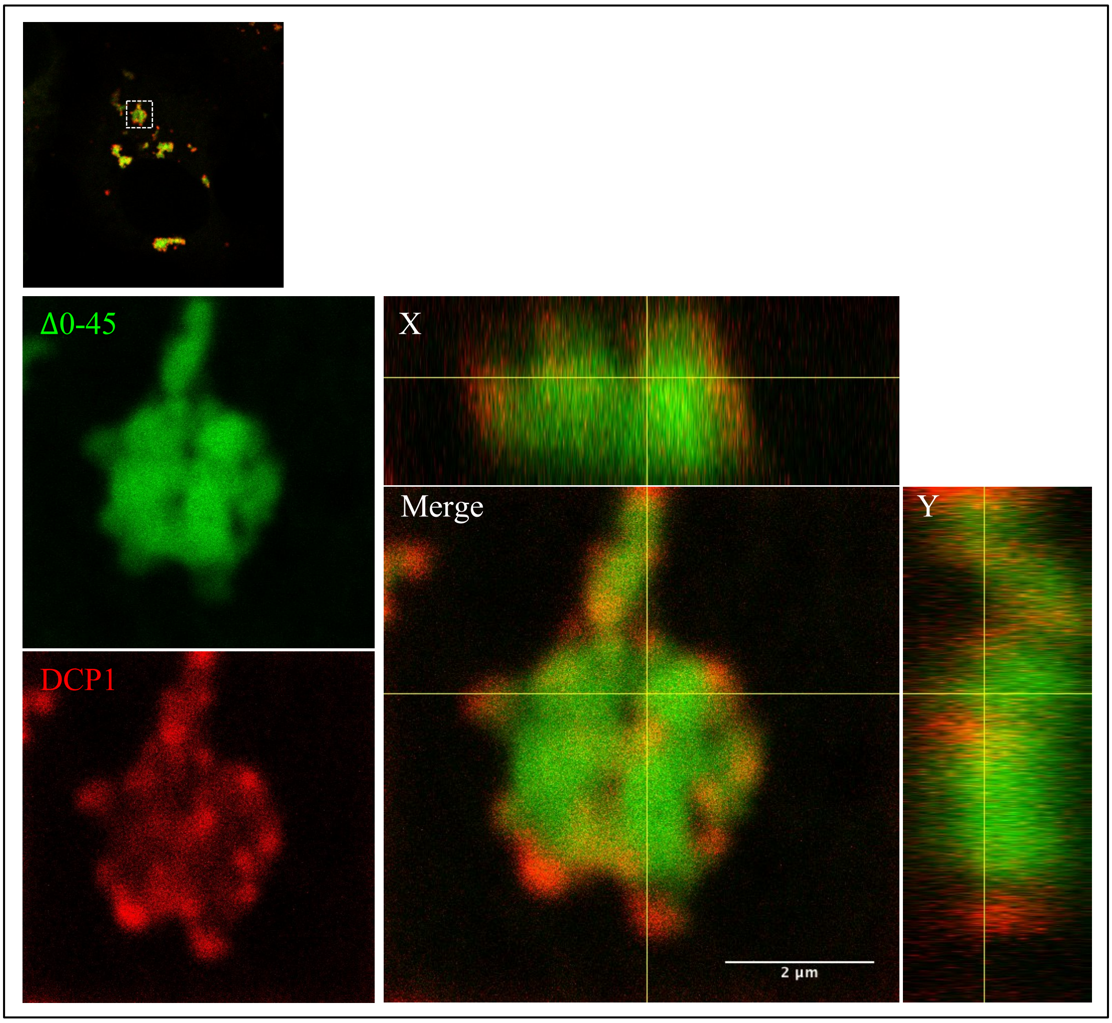


**Supplementary Figure 4.** Coiled coil-mediated aggregates (Δ0-45) of DEF6 form large structure that 'trap' DCP1. Confocal analysis of transfected COS7 cells as described before. GFP-tagged Δ0-45 formed aggregates and larger structures but in this case these structures partially overlapped with DCP1; again altering the normal cellular localisation of DCP1 (see merged images on the right).

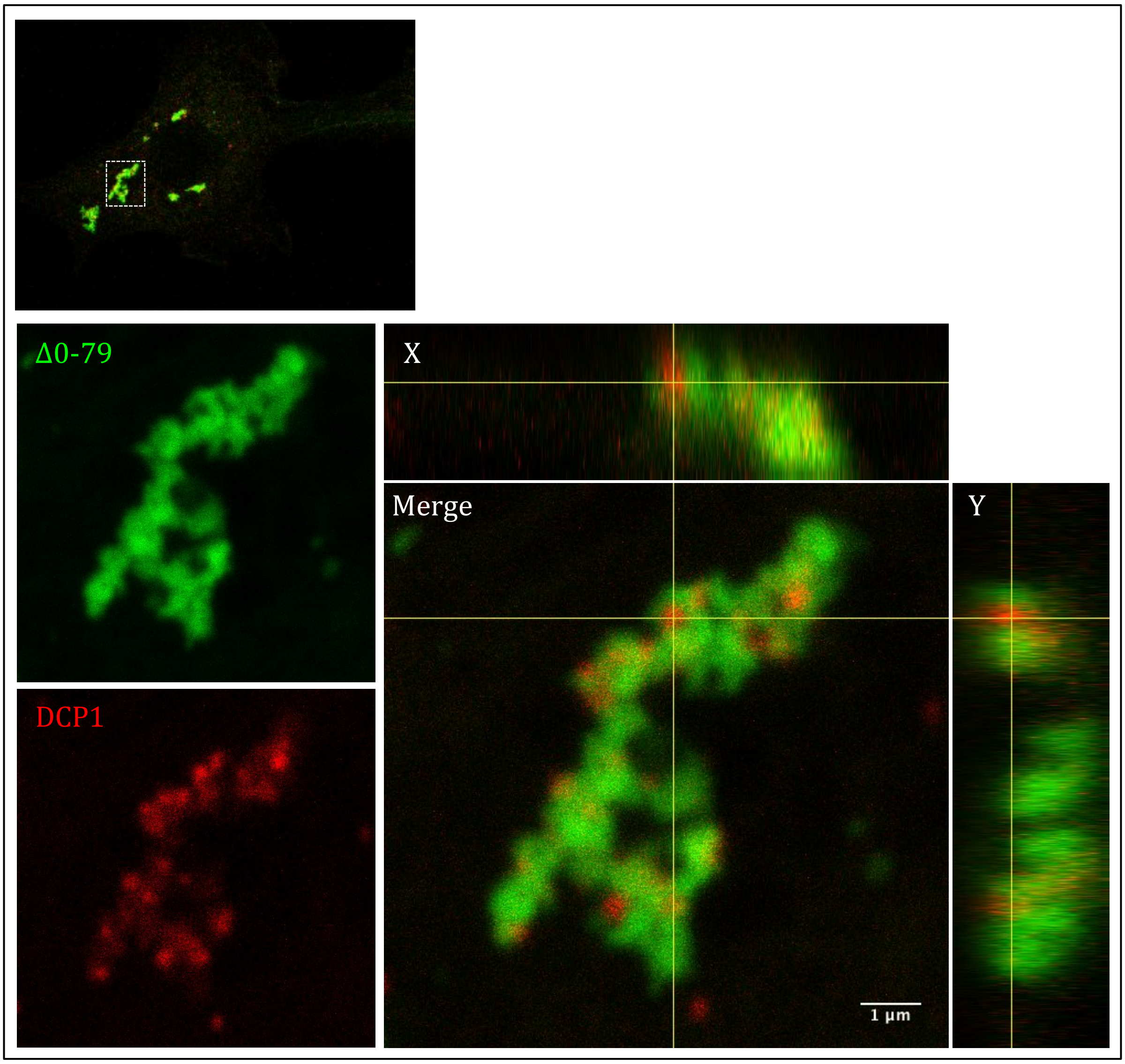


**Supplementary Figure 5.** Coiled coil-mediated aggregates (Δ0-79) of DEF6 form large structure that 'trap' DCP. Confocal analysis of transfected COS7 cells as described before. GFP-tagged Δ0-79 formed aggregates and larger structures but in this case these structures partially overlapped with DCP1; again altering the normal cellular localisation of DCP1 (see merged images on the right).

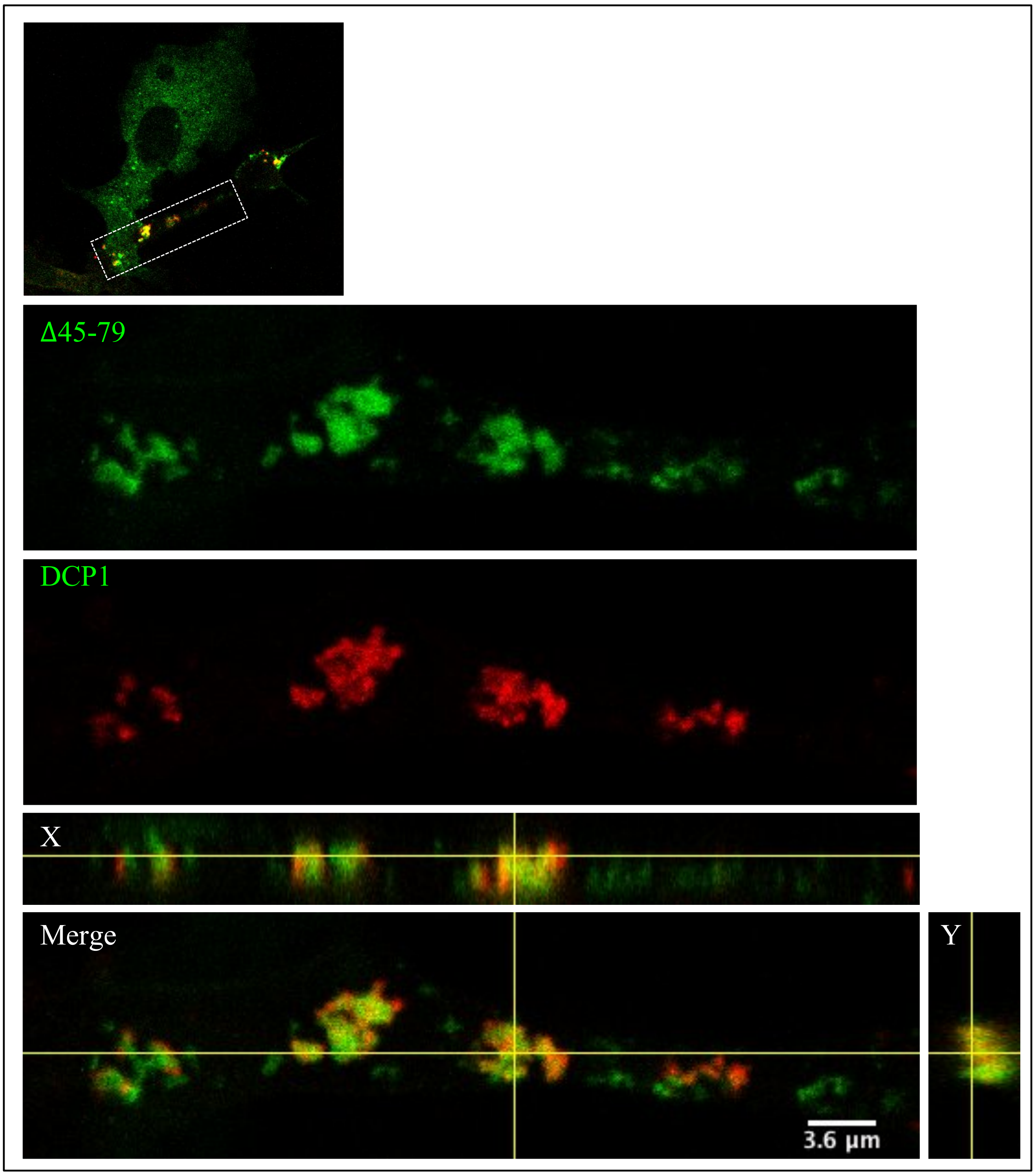


**Supplementary Figure 6.** Coiled coil-mediated aggregates (Δ45-79) of DEF6 form large structure that ‘trap’ DCP1. Confocal analysis of transfected COS7 cells as described before. GFP-tagged Δ45-79 formed aggregates and larger structures but in this case these structures partially overlapped with DCP1; again altering the normal cellular localisation of DCP1 (see merged images on the right).

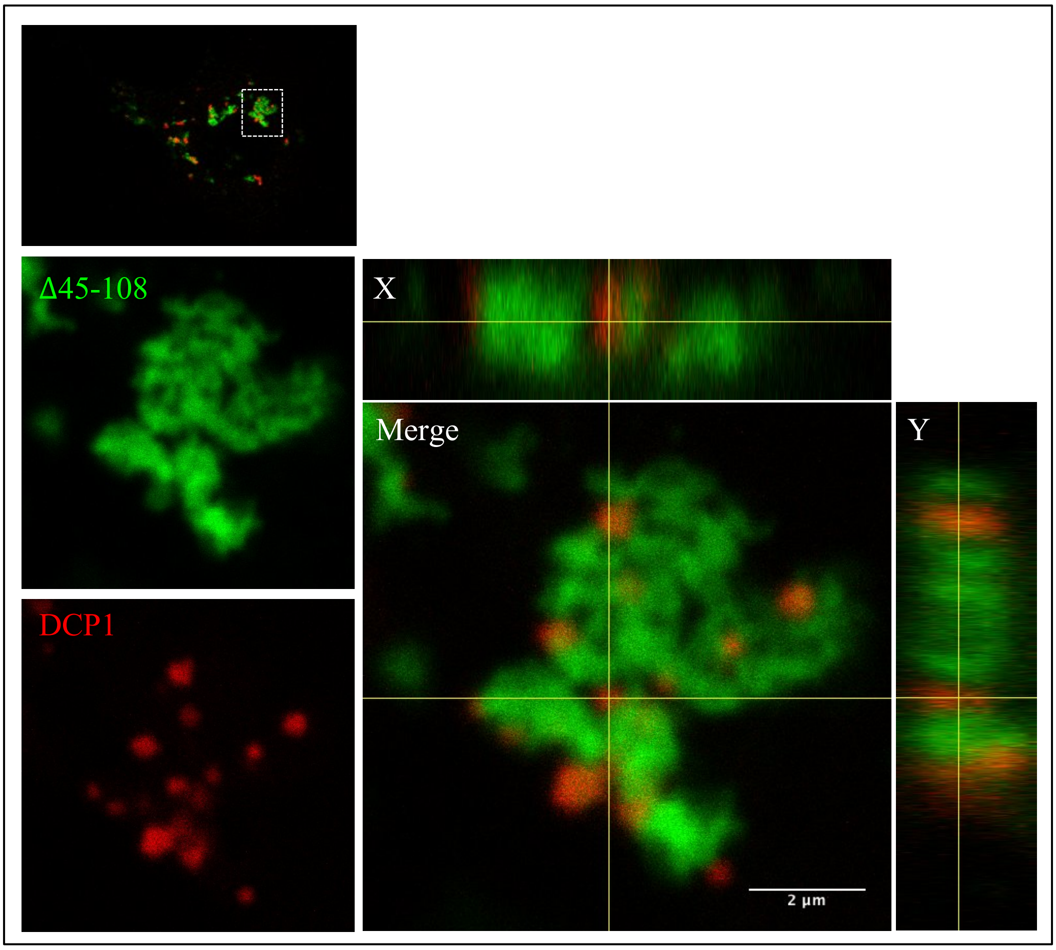


**Supplementary Figure 7.** Coiled coil-mediated aggregates (Δ45-108) of DEF6 form large structure that 'trap' DCP1. Confocal analysis of transfected COS7 cells as described before. GFP-tagged Δ45-108 formed aggregates and larger structures but in this case these structures partially overlapped with DCP1; again altering the normal cellular localisation of DCP1 (see merged images on the right).

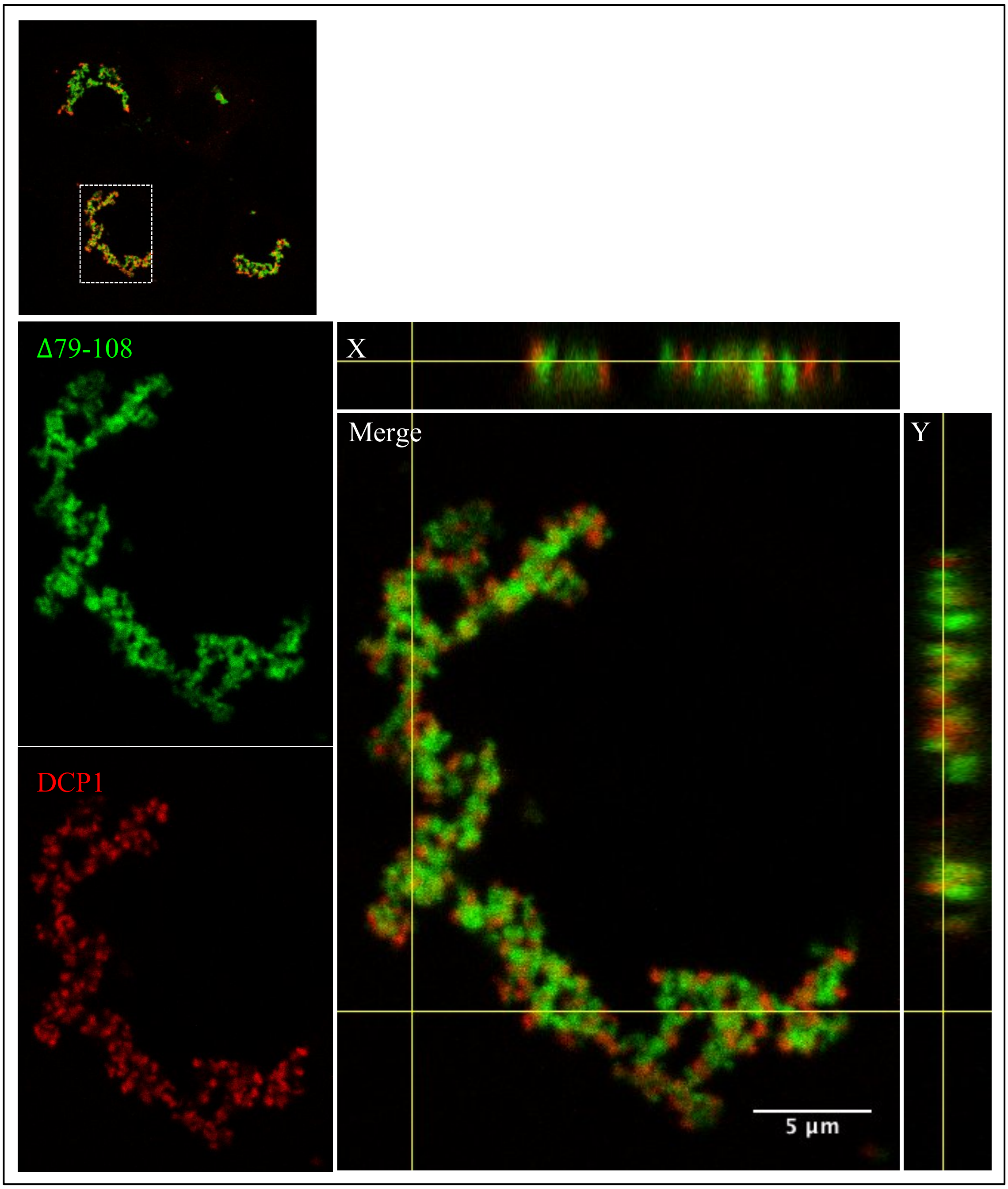


**Supplementary Figure 8.** Coiled coil-mediated aggregates (Δ79-108) of DEF6 form large structure that 'trap' DCP1. Confocal analysis of transfected COS7 cells as described before. GFP-tagged Δ79-108 formed aggregates and larger structures but in this case these structures partially overlapped with DCP1; again altering the normal cellular localisation of DCP1 (see merged images on the right).

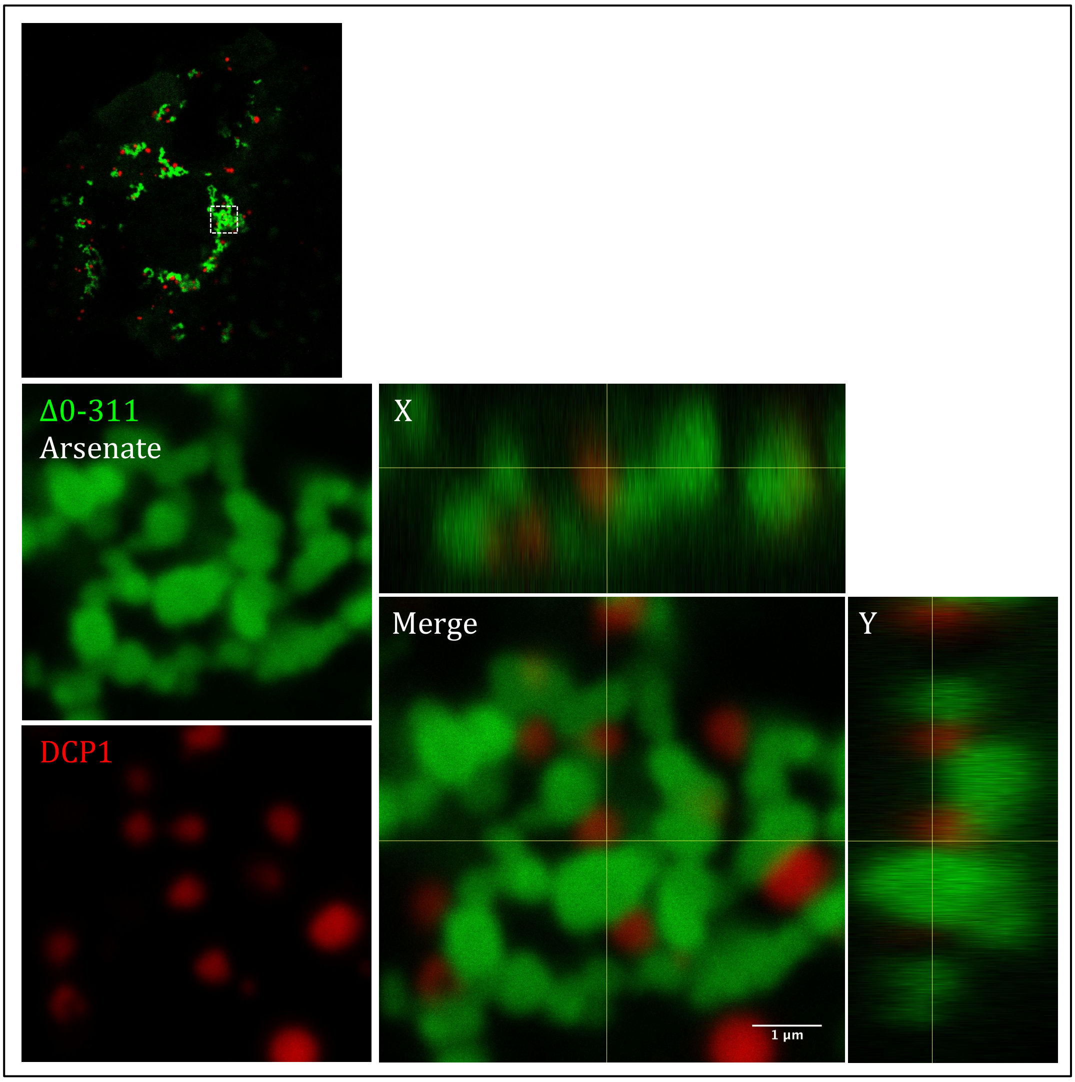


**Supplementary Figure 9.** Cellular stress (Arsenate) had no effect on formation of large structures of Δ0-311 and their ability to 'trap' DCP1. Confocal analysis of COS7 cells expressing GFP-tagged Δ0-311 and mCherry- tagged DCP1 after arsenate treatment as described before. Merged images including vertical and side views (X and Y coordinates) shown on the right indicated large structures of Δ0-311 adjacent to DCP1

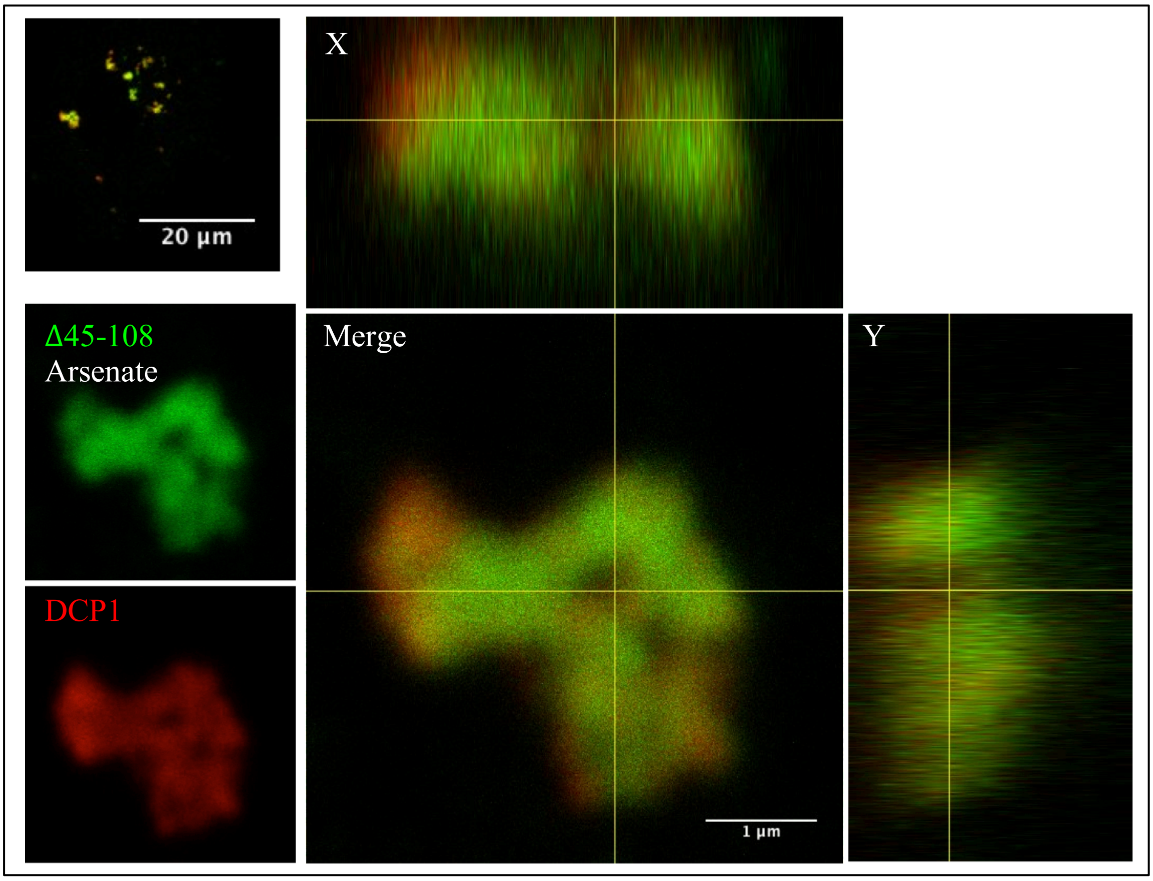


**Supplementary Figure 10.** Cellular stress (Arsenate) results in complete colocalisation of Δ45-108 structures with DCP1. Confocal analysis of COS7 cells expressing GFP-tagged Δ45-108 (representing group 3 mutants) and mCherry-tagged DCP1 after arsenate treatment as described before. Merged images including vertical and side views (X and Y coordinates) shown on the right indicated that large structures of Δ45-108 completely overlapped with DCP1.

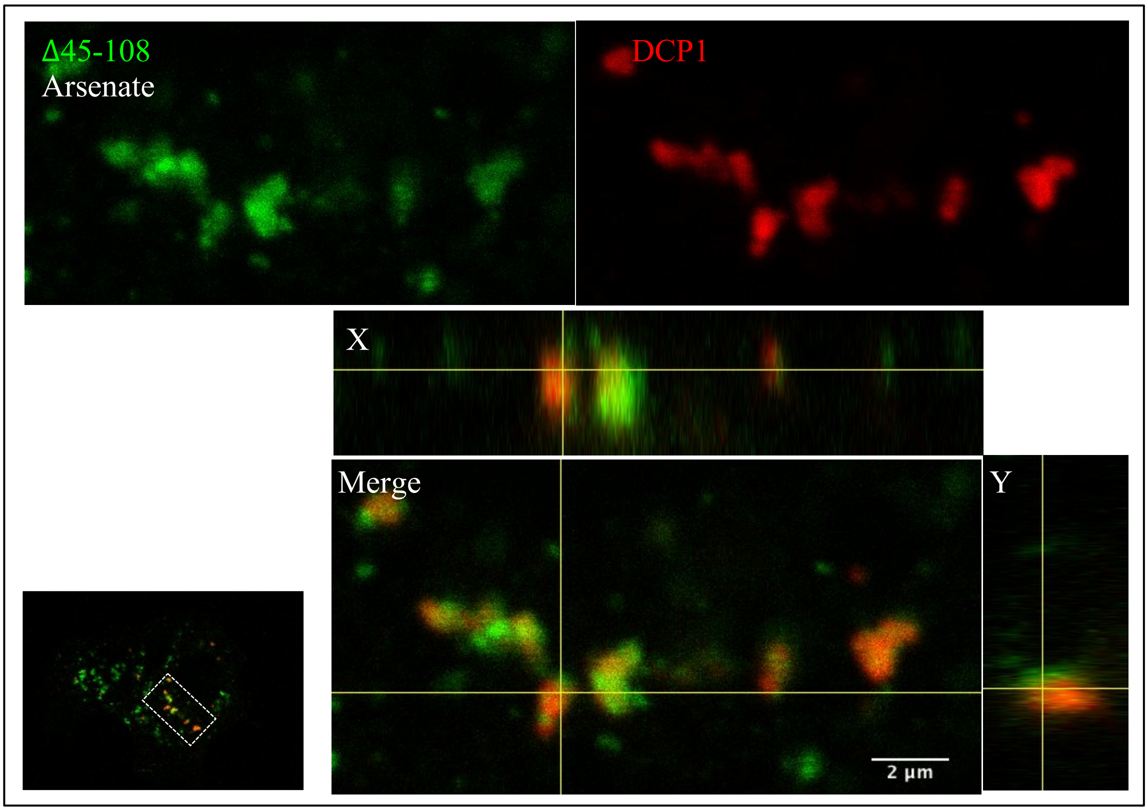


**Supplementary Figure 11.** After arsenate treatment, few samples of Δ45-108 remained in partially overlapping with DCP1. Confocal analysis of COS7 cells expressing GFP-tagged Δ45-108 (representing group 3 mutants) and mCherry-tagged DCP1 after arsenate treatment as described before. Merged images including vertical and side views (X and Y coordinates) shown on the right indicated that few large structures of Δ45-108 remained in partially overlapped with DCP1.

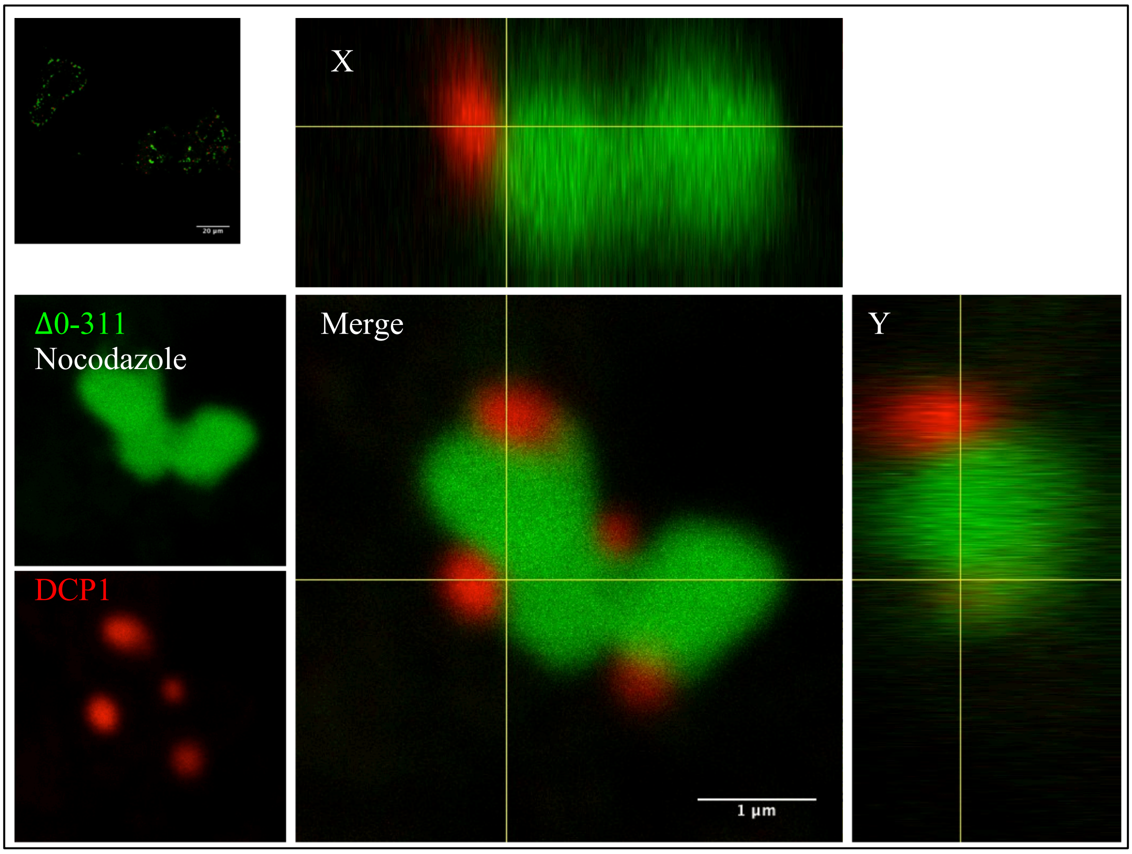


**Supplementary Figure 12.** Cellular stress (Nocodozale) had no effect on formation of large structures of Δ0-311 and their ability to 'trap' DCP1. Confocal analysis of COS7 cells expressing GFP-tagged Δ0-311 and mCherry-tagged DCP1 after arsenate treatment as described before. Merged images including vertical and side views (X and Y coordinates) shown on the right indicated large structures of Δ0-311 adjacent to DCP1

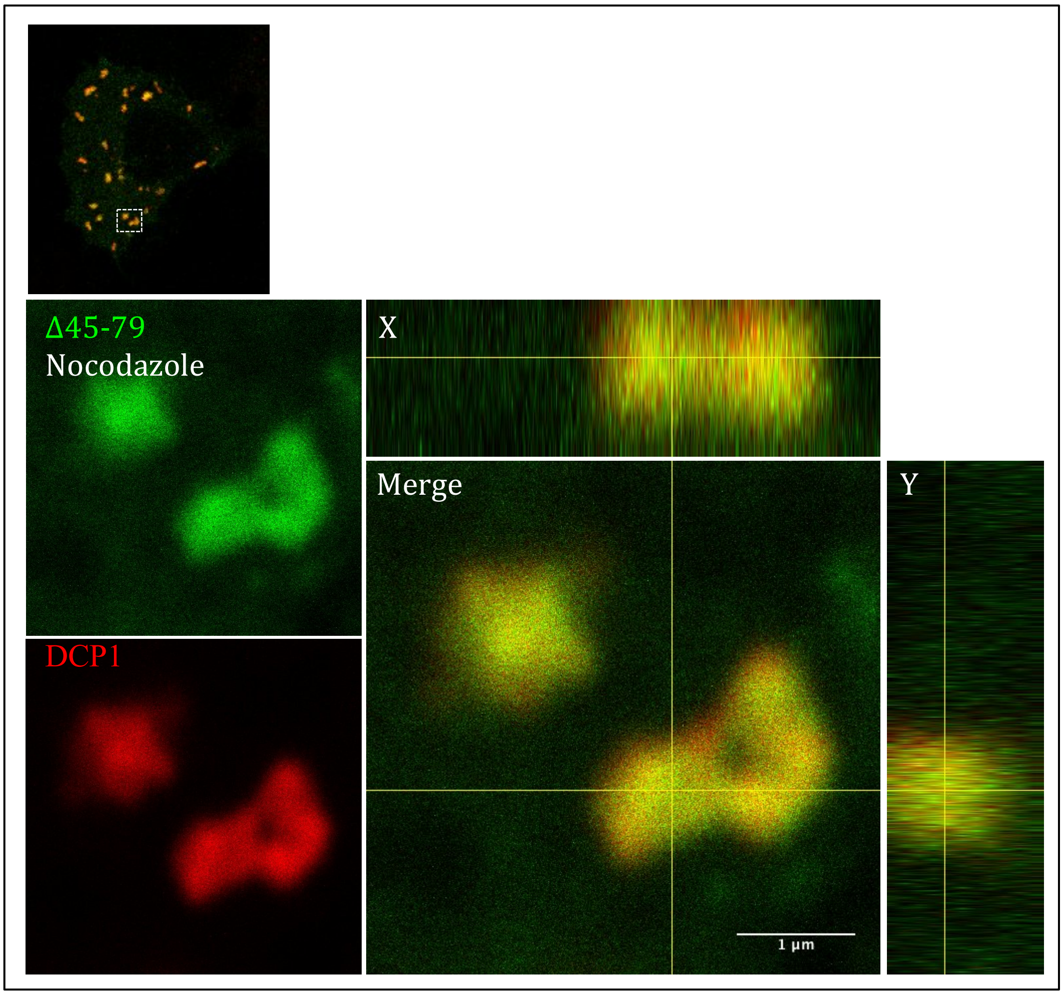


**Supplementary Figure 13.** Cellular stress (Nocodazole) results in complete colocalisation of Δ45-79 structures with DCP1. Confocal analysis of COS7 cells expressing GFP-tagged Δ45-79 (representing group 3 mutants) and mCherry-tagged DCP1 after arsenate treatment as described before. Merged images including vertical and side views (X and Y coordinates) shown on the right indicated that large structures of Δ45-79 completely overlapped with DCP1.

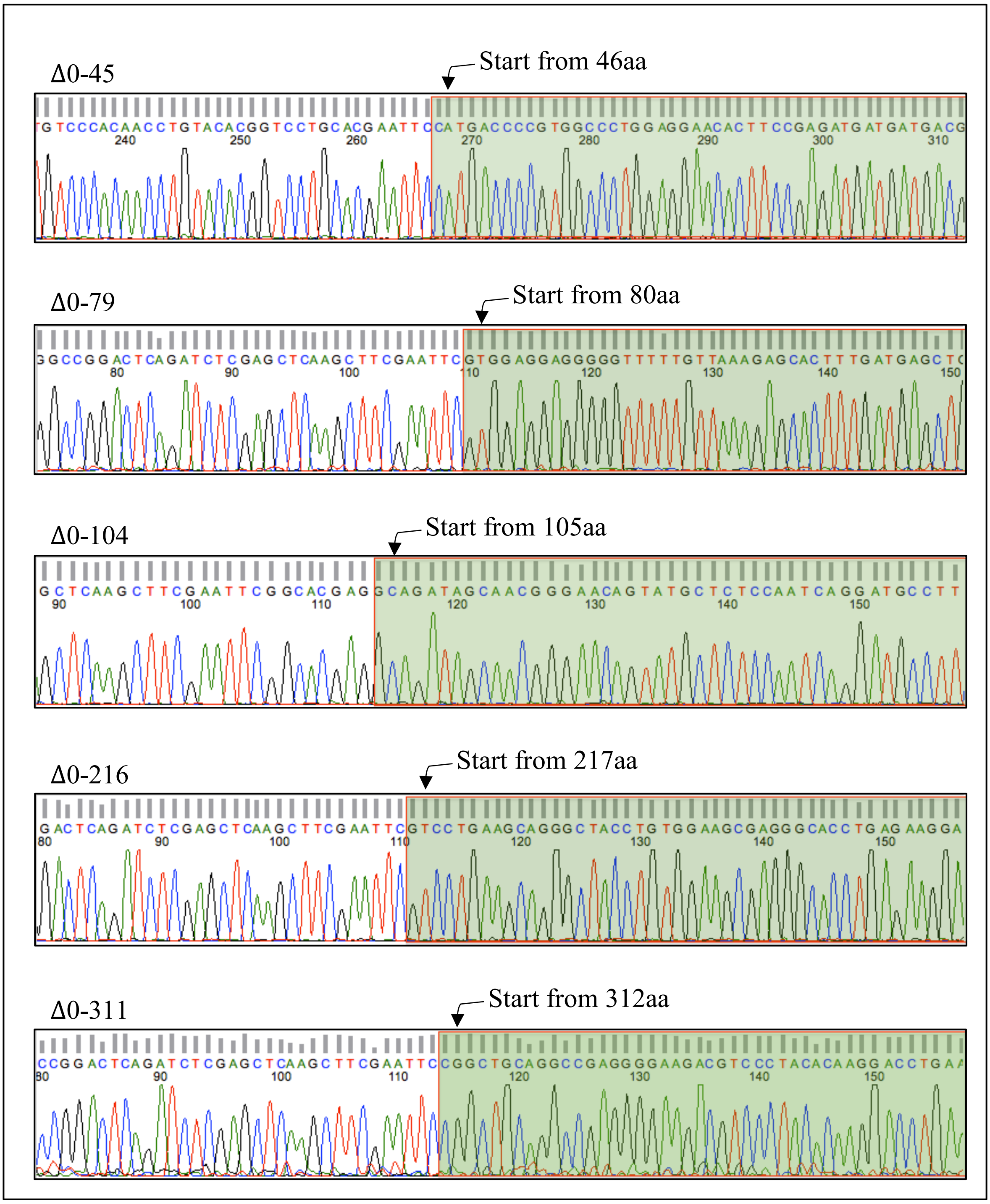


**Supplementary Figure 14.** DNA sequencing results of DEF6 N-terminal truncated mutants Green colour and arrow indicates the start amino acids of these mutants.

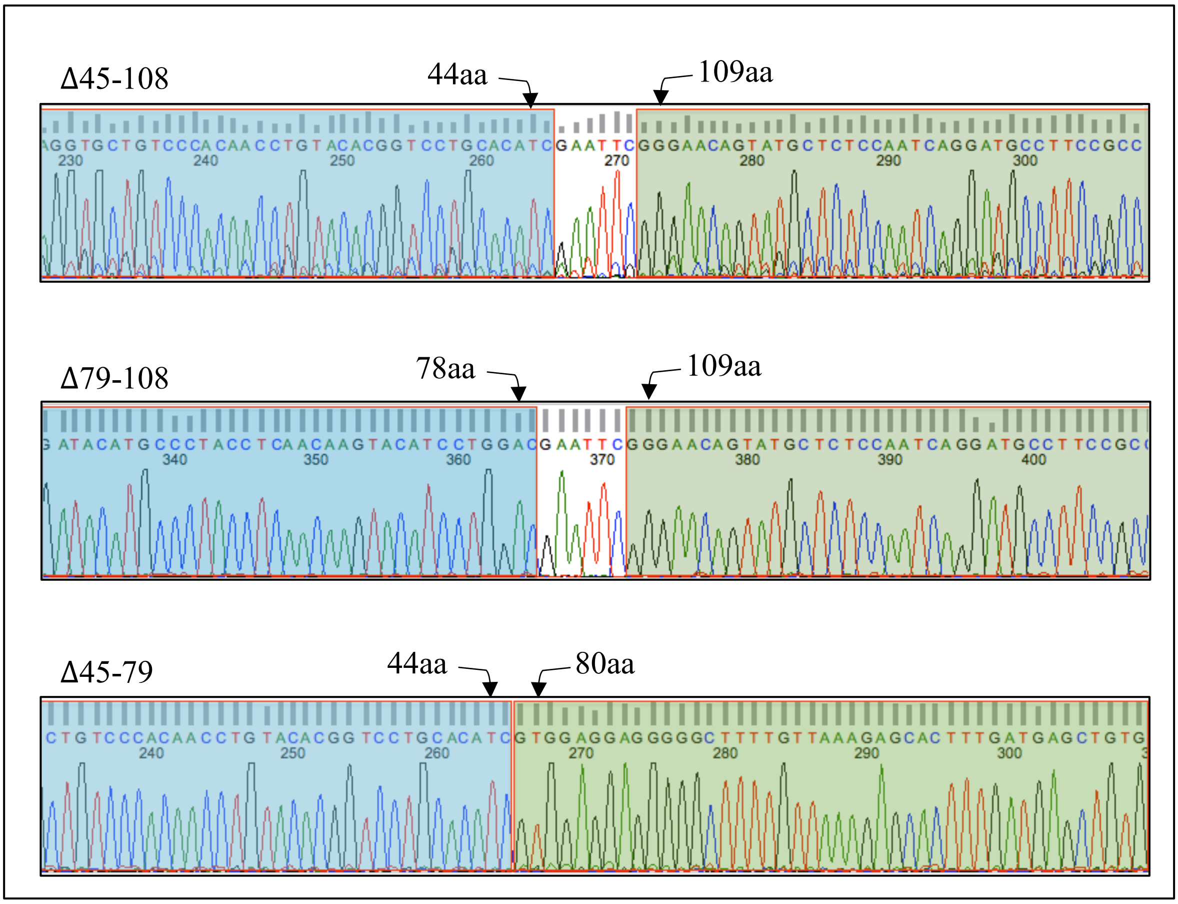


**Supplementary Figure 15.** DNA sequencing results of Δ45-108, Δ79-108 and Δ45-79. The introduced EcoRI digesting sites were remaining in Δ45-108 and Δ79-108; only Δ45-79 has been completely removed this site.

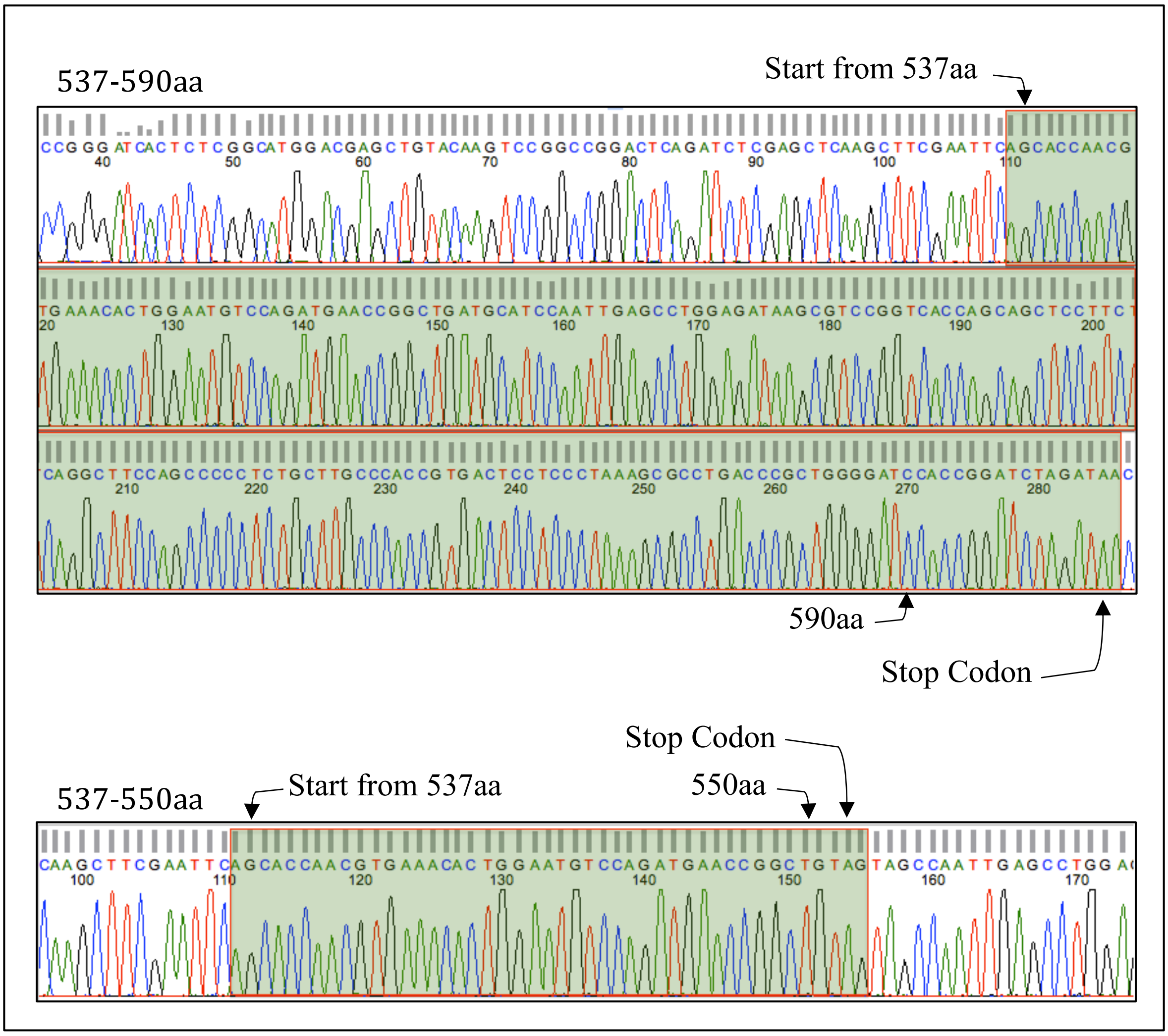


**Supplementary Figure 16a.** DNA sequencing result of DEF6 mutants 537-590aa, 537-550aa. The molecules are indicated in green with arrows to indicate start and stop positions

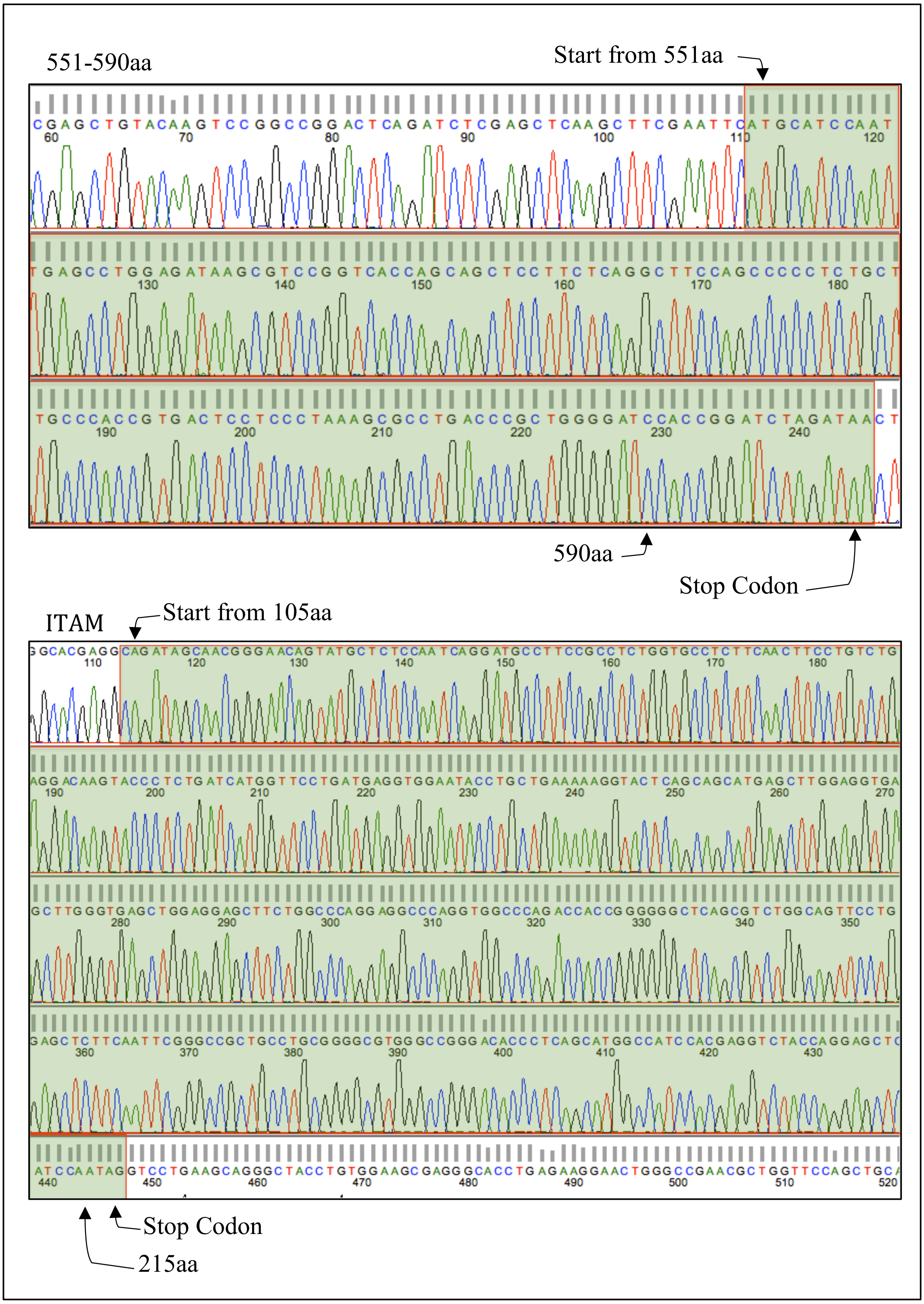


**Supplementary Figure 16b.** DNA sequencing result of DEF6 mutants 551-590aa and ITAM (105-215aa). The molecules are indicated in green with arrows to indicate start and stop positions.

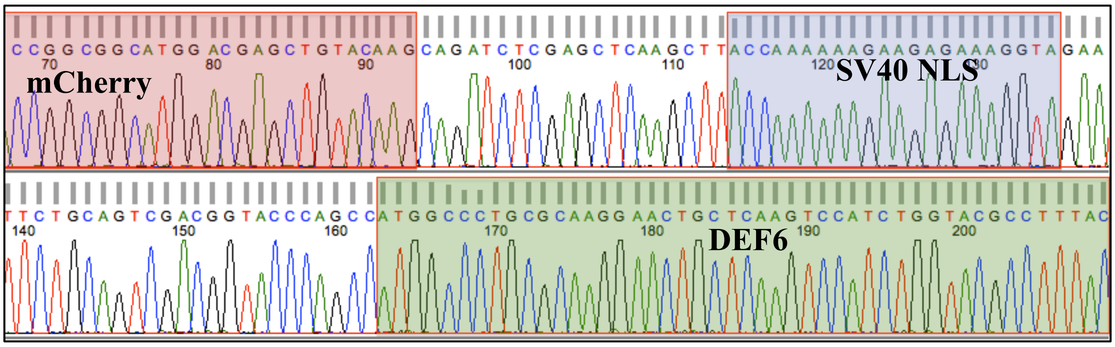


**Supplementary Figure 17.** DNA sequencing result of mCherry-NLS-DEF6. mCherry and DEF6 are respectively indicated in red and green; SV40 nuclear localization sequence (NLS) is indicated in purple that is localized between mCherry and DEF6.

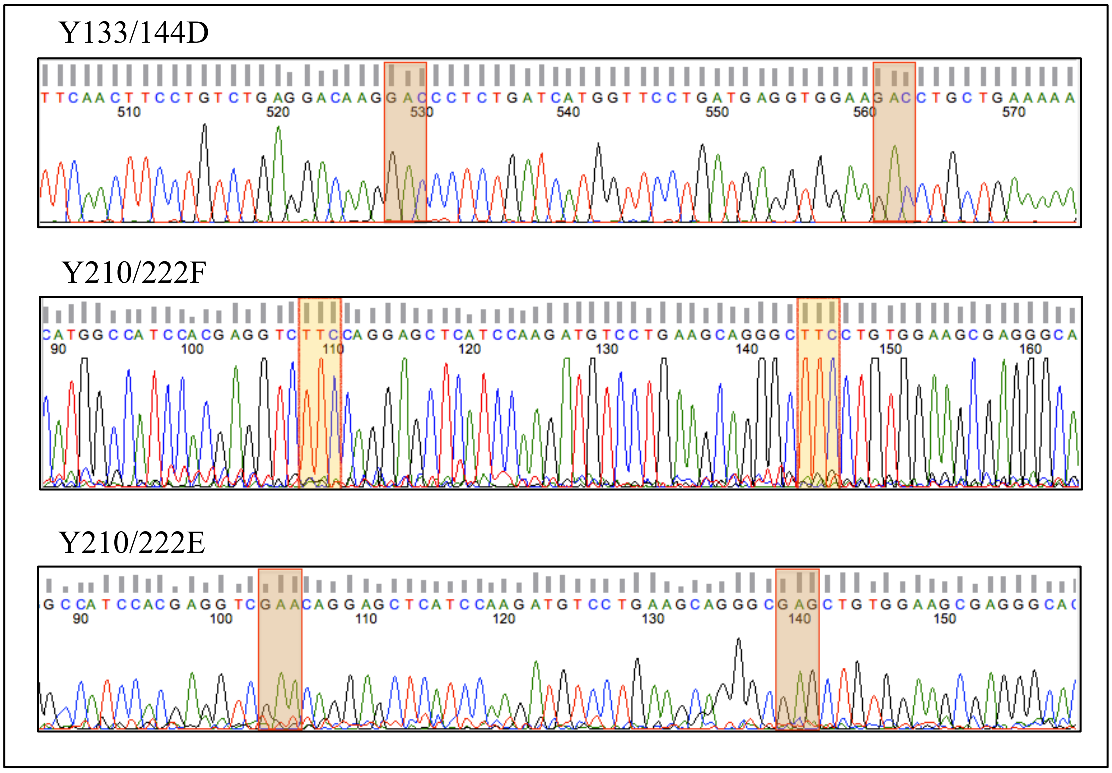


**Supplementary Figure 18**. DNA sequencing results of DEF6 phosphormimic and phosphor preventing mutants. The mutated positions are indicated by coloured boxes.
